# VH-replacement shapes the antibody repertoire by targeting non-pairing heavy-chains

**DOI:** 10.1101/2025.04.07.647564

**Authors:** Harry N White, Peter Chovanec, Laura Biggins, Elise C French, Georgia Bullen, Simon Andrews, Anne E Corcoran

**Author notes:** David Geffen School of Medicine, UCLA, CA90095, USA.

## Abstract

The diversity of antibodies underpins robust immune responses. During the formation of the antibody repertoire in early bone marrow B-cells, random antibody heavy-chain proteins are generated from recombined V, D and J gene-segments. Many are non-functional and are negatively selected. Critically, this phase of development is impacted by ageing and inflammation. To understand this process in normal mice we have undertaken the first in-depth analysis of heavy-chain selection at this ‘pre-B cell transition’. We find independent selection acting on three regions of the CDR3 antigen binding site, with particularly heavy counter-selection of certain productive VH/JH combinations. This led us to hypothesise that VH-replacement is targeting productive VDJ rearrangements that cannot pair with the surrogate light-chain (SLC). We detect VH-replacement recombination products that closely follow the pattern of counter- selection of productive VDJ. This reveals a physiological role for VH-replacement: the developmental release of B-cells that are stalled, with a productive but not SLC-pairing VDJ, leading to re-modelling of the early VDJ repertoire toward locus-distal VH that we show are more tolerant of CDR3 composition.

## Introduction

VDJ Recombination is catalysed by the RAG recombinase (Schatz et al., 1989) with D-J joining occurring first, in early pro B-cells (Alt et al., 1984). Through imprecise joining, the junction regions between D-J and V-D are diversified (Weigert et al., 1980). With the D-segments, which can be read in any translational reading frame (RF), they form the highly diverse heavy chain CDR3 regions (Figure 1a).

**Figure 1.**
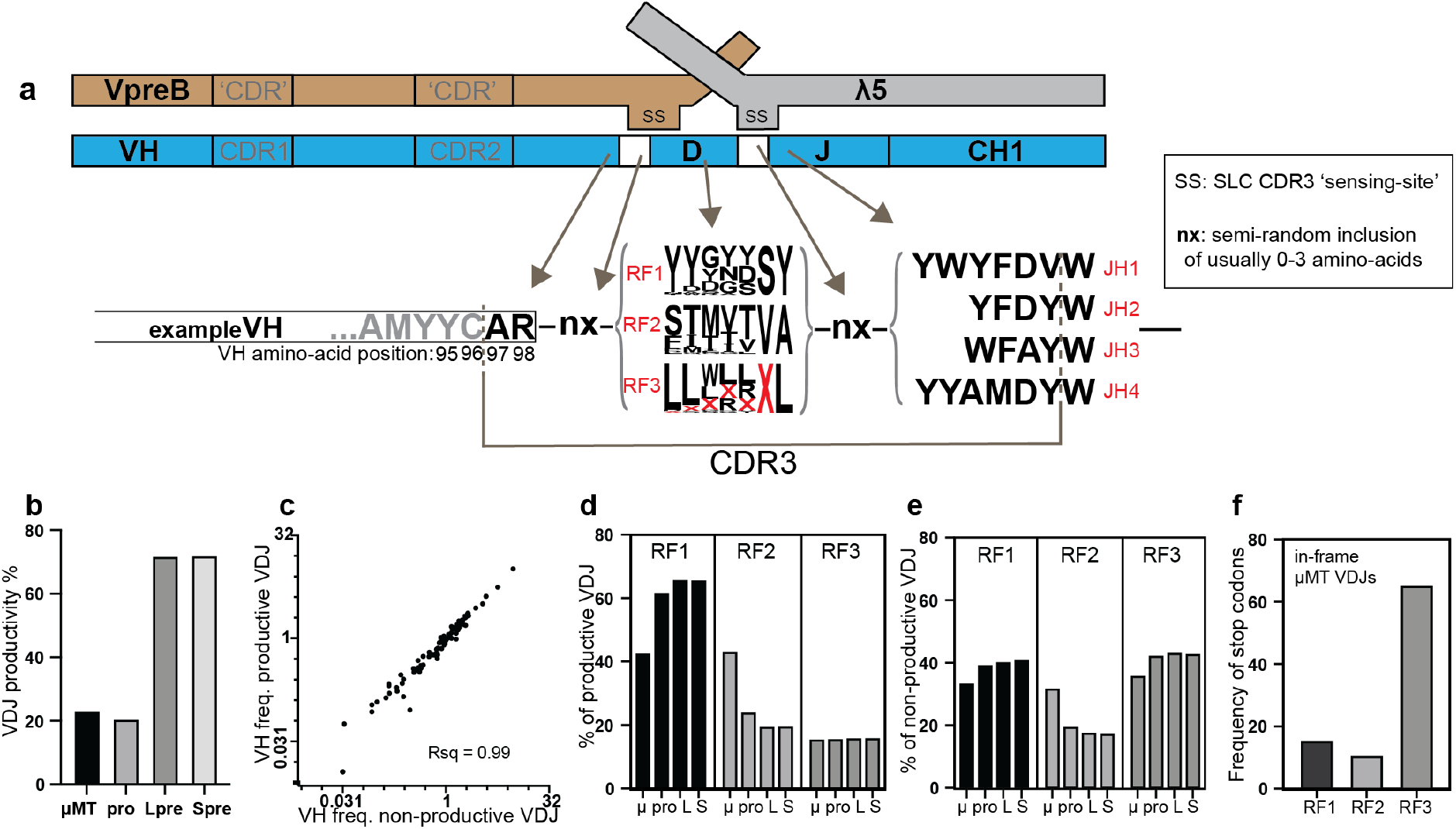
CDR3 sequence structure and overall VDJ productivity, reading-frames and stop-codons. **a**, Amino acid sequence structure of the heavy-chain CDR3 showing contribution of different gene segments and random elements to CDR3 sequence. We use the convention that the invariant Cysteine at the 3’ end of the VH framework-3 region (FR3) is residue 96 and the CDR3 begins at residue 97 and ends before the conserved Tryptophan (W) residue in JH. The logo plot of frequencies of amino-acid use in the different D reading frames, RF1-3, was calculated using D-region sequences factored by their frequency of use in mMT pro-B cells. The red X represents a stop codon. The regions nX usually result in the net inclusion of 0 to 3 amino-acids. This outcome is the result of inclusion of palindromic and non-templated nucleotides by Artemis hairpin-cleavage and TdT activity respectively, and also exonuclease removal of nucleotides, all during VDJ recombination. Thus, the terminal residues of the VH, D and JH can also be altered. **b**, Frequency of productive VDJ formation in the four cell types analysed: mMT, mMT pro-B cells; pro, pro-B cells; Lpre, large pre-B cells; Spre, small pre-B cells. **c**, Frequency of individual VH in non-productive and productive VDJ from mMT pro-B cells. **d**, Relative frequency of different D- reading frames in productive VDJ from the four cell-types analysed:, mMT pro-B cells; pro, pro-B cells; L, large pre-B cells; S, small pre-B cells. **e**, Relative frequency of different D-reading frames in productive VDJ from the four cell-types analysed: mMT pro-B cells; pro, pro-B cells; L, large pre-B cells; S, small pre-B cells. **f**, Frequency of stop codons in in-frame mMT VDJ by reading-frame.

A first selection occurs against recombinants using RF2 and RF3. In-frame with most D-regions in RF2 in mouse B cells is an ATG start codon, leading to expression of a truncated ‘Dμ’ protein and developmental arrest (Tornberg et al., 1998). Many D-regions contain stop codons in RF3 (Figure 1a). Human B-cells use other processes to select RFs as Dμ-selection doesn’t occur (Minegishi & Conley, 2001).

VDJ recombinants that are in-frame without stop codons, ‘productive’, are expressed as heavy chain – ‘μ’ protein. Co-expressed with μ is the SLC, formed from two components, VpreB and λ5, which may pair with μ to form the pre-B cell receptor (pre-BCR). Successful pre-BCR signalling from a ‘functional’ pairing heavy-chain, results in suppression of SLC expression (Grawunder et al., 1995), allelic exclusion, and several cell divisions, finishing the pre-B cell transition (Martensson et al., 2010).

Importantly, as many as half of new μ-chains cannot pair with the SLC, and pairing efficiency differs between VH families (ten Boekel et al., 1997), although sample numbers in this study were low.

A pre-BCR structure shows that particular amino-acid residues of the SLC make extensive contact with residues that bracket the CDR3H, forming a ‘sensing site’ (Bankovich et al., 2007). Thus, the variable stability of μ-chain/SLC pairing, impacting the longevity of the pre-BCR (Lassoued et al., 1996), would drive differential VDJ selection over the pre-B transition, (Malynn et al., 1990).

Poor μ-chain pairing with the SLC could be VH-intrinsic, or depend on the CDR3. Specific instances of CDR3 sequence selection over the pre-B transition have been reported, supporting the latter. Those few VH5-2/81X heavy chains that survive the pre-B transition often use Histidine at position 99 in the CDR3H (Martin et al., 2003).

Similarly, selection for tyrosine residues in VH position 101 in VH5/7183 genes occurs (Khass et al., 2016).

An additional process that may contribute to repertoire alteration is VH-replacement (VHR). Here, a pre-existing VDJ is invaded by an upstream VH on the same chromosome, that replaces the original VH creating a new VDJ (Koralov et al., 2006; Usuda et al., 1992). Most VH have a ‘cryptic’ 7-mer recombination-signal-sequence (cRSS) close to their 3’ end that could facilitate this. The impact of VHR on VDJ repertoire organisation is unknown. It was thought to enable heavy-chain receptor-editing (Chen et al., 1995) but this was disproven (Sun et al., 2015). VHR occurs in pro-B cells and is Rag-dependent (Cowell et al., 2003; Davila et al., 2007). In a mouse carrying a monoclonal functional VDJ, subsequent VHR can contribute up to 20% of the peripheral antibody repertoire (Kumar et al., 2015).

## Results

We undertook the first in-depth analysis of VDJ selection over the pre-B transition. Sequence metadata are in table S1a. μMT mice were also analysed; the μMT mouse has a deletion in the IgM transmembrane domain abrogating signalling and causing a developmental block at the pro-B cell stage (Kitamura et al., 1991). We analysed the 89 most frequently used VH in VDJ recombinants from pro-B cells, that form 98% of the repertoire.

### Less than a quarter of heavy-chain VDJ rearrangements are productive

We found that only 23% of VDJ recombinants are productive in μMT mouse pro B-cells (Figure 1b and table S1). High correlation between productive and non-productive VH gene frequency in μMT pro B-cell VDJ confirms this figure is not impacted by selection (Figure 1c), thus establishing it as intrinsic to mouse VDJ recombination.

### D reading-frame selection halves the frequency of VDJ with RF2 and RF3

Dμ-selection, that does not occur in μMT mice (Gu et al., 1991), halves the relative frequency of RF2 VDJs (Figure 1d and 1e). Stop codons more than halve productive rearrangements in RF3 (Figure 1d). No D-regions have a stop codon in RF2, in the overwhelmingly favoured forward orientation. Thus, the 10% of RF2 in-frame VDJ with a stop codon indicates the frequency stop codons are formed *de novo* during VDJ recombination (Figure 1f).

### Overall VH selection from pro-B to pre-B cells

Importantly, most productive VDJ from pro B-cells show selection prior to pre-BCR mediated proliferation (Figure 2a/S2a). We have used non-productive VDJ from pro B-cells, therefore, as a measure of un-selected VDJ rearrangements. Figure 2b shows the VH frequency from these versus the productive VDJ repertoire from small pre-B-cells, by locus position. This indicates overall VH selection over the pre-B transition. As well as the most proximal gene, VH5-2 (81X), which shows a large drop from 12.3% of total repertoire to 0.5%, the trend is for the more frequently rearranged proximal VH to be strongly counter-selected, and the more distal VH to be positively selected, confirming previous reports for small numbers of VH genes (Malynn et al., 1990), although VH3- 1,12-3 and 1-11 do not follow the trend.

**Figure 2.**
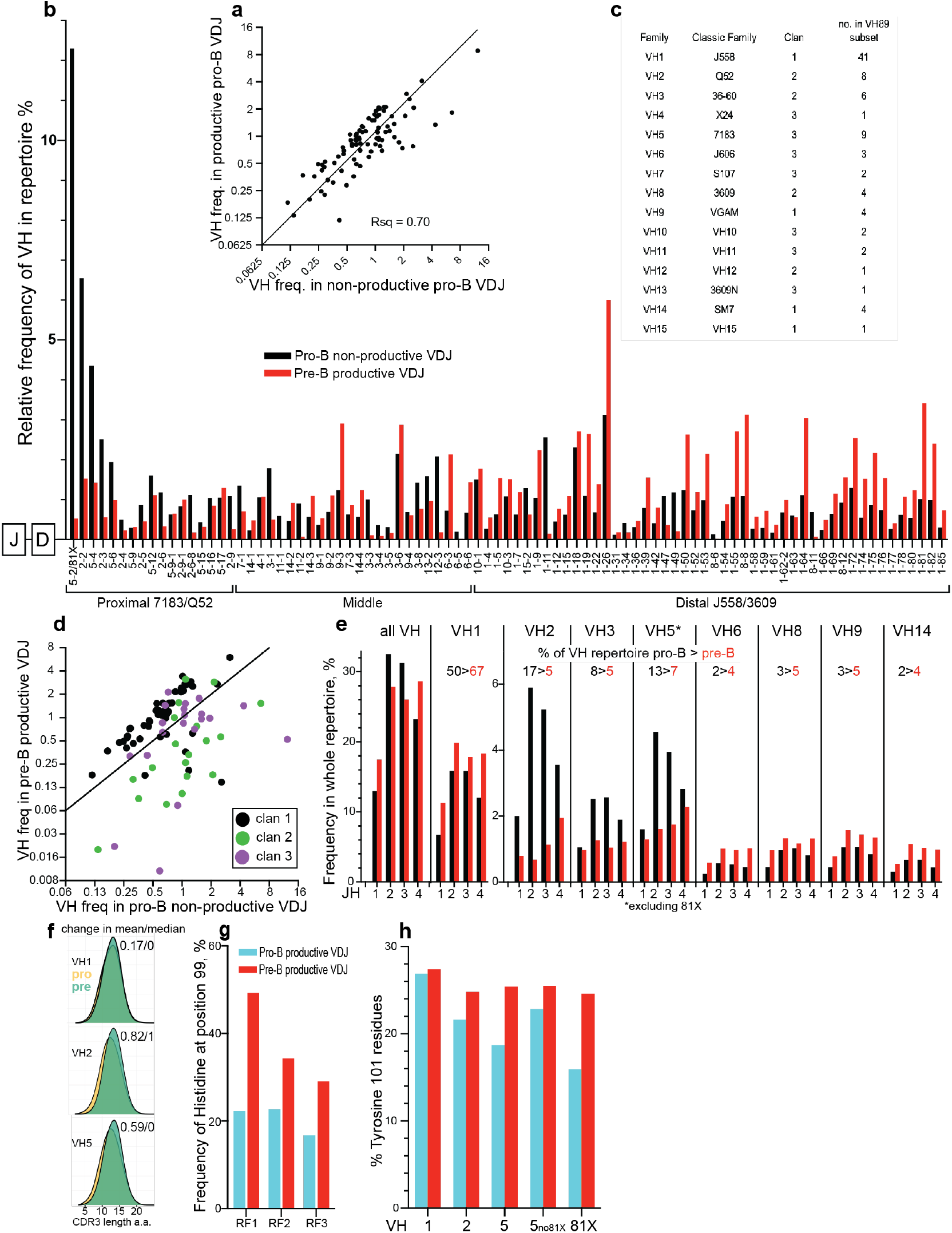
Selection of VH genes and families, JH and particular CDR3 amino-acids, in the VDJ repertoire over the pre-B cell transition. **a**, Frequency of individual VH in productive and non-productive VDJ from pro-B cells. **b**, Frequency of individual VH by IgH locus position, over the pre-B cell transition, as measured from non-productive VDJ from pro-B cells and productive VDJ from small pre-B cells. **c**, Table showing VH gene families, their classic names, clans, and the number of genes in the family as detected in the group of 89 most used VH. **d**, plot of frequency of individual VH in pro-B cell non-productive VDJ versus small pre-B cell productive VDJ, coloured by VH clan. **e**, Frequency of particular VH-family/JH combinations as a proportion of the entire VDJ repertoire from pro-B cell non-productive VDJ and small pre-B cell productive VDJ. Numbers above bar plots refer to the overall frequency of that VH family in the VDJ repertoires measured. Note plots for VH5 exclude data from VH5-2/81X. **f**, CDR3 length distribution for VH1, VH2 and VH5 families over the pre-B transition: pro, pro-B cells; pre, small pre-B cells. Numbers refer to the difference in the mean/median CDR3 length between these cell populations. Data from RF1 sequences only. **g**, Frequency of Histidine residues at VH5-2/81X VDJ position 99 in the pro-B cell and pre-B cell productive VDJ repertoires, by D-reading frame. **h**, Frequency of Tyrosine residues at VDJ positions 101 in the pro-B cell and pre-B cell productive VDJ repertoires, by VH-family. 5no81X, data for VH5 family excluding data for VH5-2/81X; 81X, data for VH5- 2/81X alone.

The overall VH selection shown in Figure 2b is the net outcome of 2 stages of change detected by our methods. We detect selection between non-productive and productive VDJ in pro-B cells (Figure S2a), and further selection between productive VDJ from pro- B to pre-B cells (Figure S2b).

VH genes can be sub-divided into three clans defined by structural features conserved in evolution (Kirkham et al., 1992). Figure 2d shows that most clan 1 VH are positively selected at the pre-B transition by around 2-fold; 12/19 clan 2 VH show a greater than two-fold counter selection; and clan 3 VH are more variably impacted.

### Strong VH-specific JH selection over the pre-B transition

JH regions are good proxies for the C-terminal part of the CDR3/junction region, as the sequence they can contribute varies in length and composition (Figure 1a). Figure 2e shows the change in frequencies of the eight largest VH families, per JH, over the pre-B transition. All VH families that show increased frequency show little differential JH selection. The three counter-selected families (VH2/3/5) show strong JH2 and JH3 negative selection and less or no selection against JH1 and JH4. JH2 and JH3 are two amino-acids shorter in their CDR3 contribution (Figure1a), their counter-selection resulting in increased length CDR3s (Figure 2f). We suggest the extra tyrosine residues in JH1 and JH4 are closer to the λ5 sensing site providing the necessary stability for VH2/5 pre-BCR formation (Bankovich et al., 2007).

This surprisingly strong VH-family specific JH selection reflects most of the negative VDJ-selection occurring at the pre-B transition. It results in removal of 21% of the repertoire, mostly JH2 and JH3 bearing proximal VH2/3/5. This contrasts with the distal VH1s which, at the level of JH, show no restriction in CDR3 diversity over the pre-B transition.

### Selection of key CDR3 amino-acids over the pre-B transition

To allow direct comparisons between CDR3 amino acid sequences from pro-B and small pre-B-cells, we have compared productive VDJ from each dataset. Figure 2g confirms selection for H99 in VH5-2/81X VDJ, also showing that H99 selection occurs for VDJ in all RFs. Figure 2h shows Y101 analysis for the 3 largest VH families. VH5 shows the greatest Y101 selection, although this appears mostly due to strong counter- selection of VH 5-2/81X VDJ, that have lower levels of Y101. Excluding VH5-2/81x, the VH2 family shows the strongest selection for Y101.

Figure 3a shows overall changes in amino acid composition of the N-terminal of CDR3s over the pre-B transition. A97 and R98, already very common in V-D junctions, are further selected in all the major VH family VDJs shown. They are VH encoded, and their selection is CDR3-independent as it occurs in all CDR3 reading-frames (Figure S3a), underlining their importance in these positions for functional CDR3s. The strong selection of A97 in VH1s follows the counter-selection of VH1-11, the only VH1 with a G97. VH1-11 falls from 2.1% to 0.15% of the total VH repertoire over the pre-B transition (Figure 2b). H99 is selected in other VH5, in addition to VH5-2/81X, but counter selected in VH2-family VDJ (Figure 3a). The counter-selection of S/K98 in VH2 genes suggests, as for G97 in VH1-11, that it may be these N-terminal CDR3 amino-acids, as well as perhaps other VH attributes, that inhibit pre-B transition of such VH.

**Figure 3.**
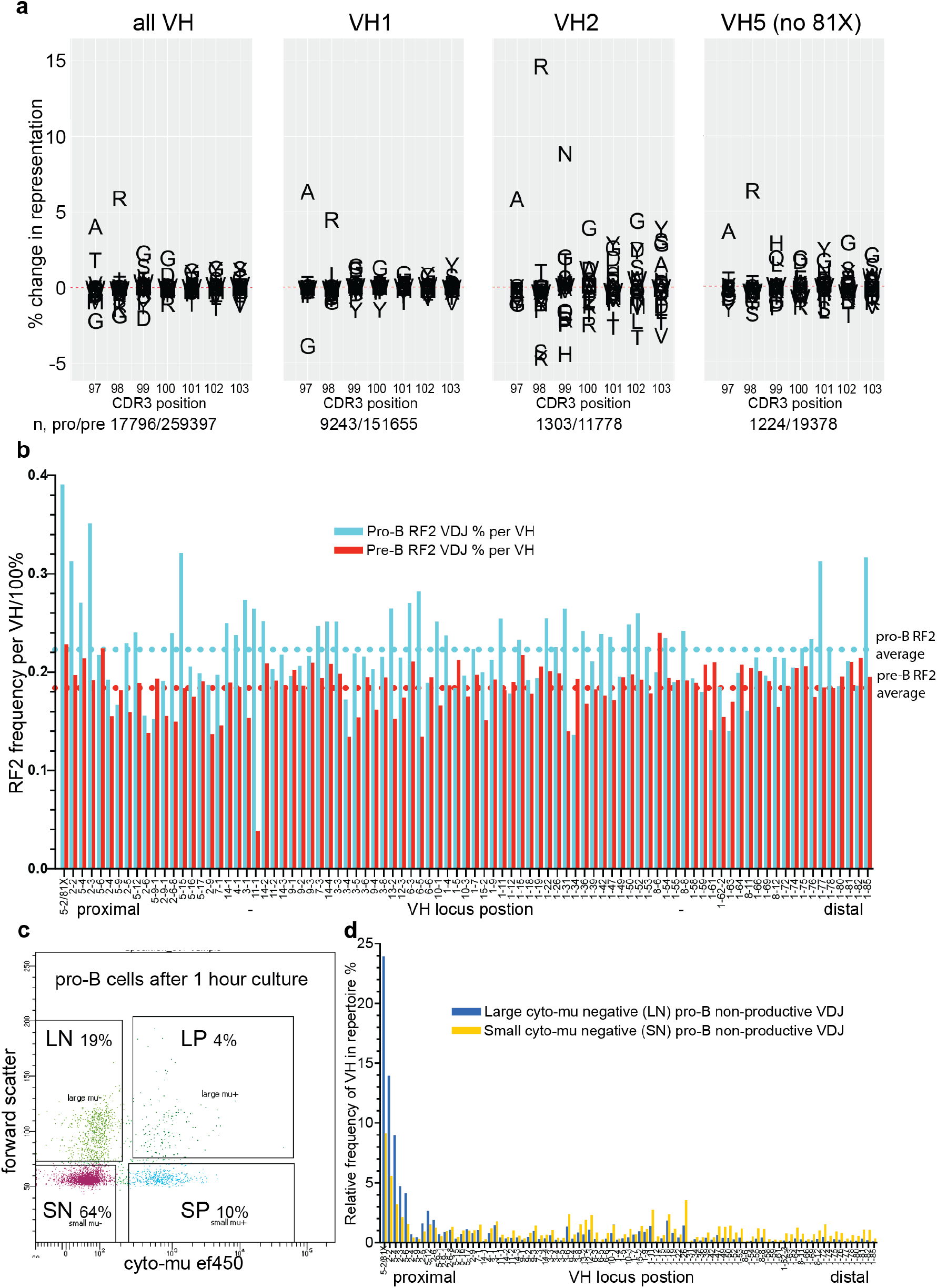
Change in amino-acid frequency by CDR3 position, and RF2 counter-selection, over the pre-B cell transition. **a**, Change in frequency of amino-acid residue use for the three major VH families, in the first seven positions of the CDR3, over the pre-B transition as measured between pro-B cell and small pre-B cell productive VDJ. Numbers below panels show numbers of sequences analysed for each VH family from pro- and pre-B cell VDJ repertoires. **b**, Frequency of RF2 D-reading frame per VH over the pre-B transition, plotted by locus position, as measured between pro-B cell and small pre-B cell productive VDJ. **c**, FACS plot showing separation and gating for sub-setting pro-B cells, after one hour of culture, into four populations based on expression of cytoplasmic-chain and forward scatter. LN, large-chain negative cells; SN, small-chain negative cells; SP, small –chain positive cells; LP, large-chain positive cells. **d**, Frequency of individual VH in non-productive VDJ repertoires from large-chain negative (LN) pro-B cells and small-chain negative (SN) pro-B cells, by locus position.

### Little selection for CDR3 charge, hydropathy and aliphatic index over the pre-B transition

Arginine (R) residues in the CDR3 were thought to be counter-selected over the pre-B transition (Keenan et al., 2008). We find no evidence for this (Figure S3b/f). Further, we find little change in the overall CDR3 biophysical properties (Figures S3c-e/g-i).

### Variable Dμ selection is followed by efficient RF2 counter-selection at the pre-B transition

Figure 3a also shows that from position 100 onwards, G and Y appear to be positively selected and T and V are counter-selected, particularly for VH2 and VH5. This is consistent with RF selection, as G/Y are common in RF1 and T/V are common in RF2 (Figure 1a). RF2 counter-selection occurs prior to the pre-B transition (Figure 1d), (Kitamura et al., 1991; Zemlin et al., 2008), although in mice expressing excess levels of RF2 VDJ, the pre-B transition is negatively impacted (Khass et al., 2016), suggesting RF2 counter selection may also occur at this later point.

The selection of RF2, per VH, over the pre-B transition is shown in Figure 3b. Whilst the average change is −4% there are some striking differences in the frequency of RF2 VDJ. 39% of VH5-2 VDJs are RF2, similar to the overall 43% RF2 in μMT pro-B cells (Figure 1d). Other proximal VH (VH2 and VH5) also show high RF2 proportions as do two distal VH1. All of this ‘excess RF2’ is eliminated at the pre-B transition, demonstrating that whether or not Dμ-mediated RF2 counter-selection has occurred, further selection at the pre-B transition is sufficient to reduce RF2 use. This explains how RF selection can occur in humans, where Dμ-selection is absent (Benichou et al., 2013; Minegishi & Conley, 2001).

We hypothesised that proximal VH recombine before Dμ-selection has had time to manifest. After one hour of culture, pro-B cells can be subset into four populations based on intracellular IgM and size (Figure 3c). One of these populations, large cyto-μ- negative cells (LN) are the least mature pro-B cells as they show the lowest frequency of V-D recombination (Supplementary Table 1a). Figure 3d shows the VH frequency of these compared to the small cyto-μ negative cells (SN) that represent the bulk of pro- Bs. These LN pro-B cells show a strong bias of V-D recombination to the most proximal VH, with VH 5-2 making up 24% of the total VDJ, and the most proximal 4 VH accounting for 52% of VDJs. Dμ levels need to be high to stop V-DJ recombination (Reth et al., 1985), and clearly the most proximal VH are recombining before Dμ inhibition has been established, explaining their high RF2 levels (Figure 3b).

### Clan-specific VDJ selection prior to pre-BCR mediated proliferation

We have shown differential selection of VH prior to pre-BCR mediated proliferation (Figures 2a, S2a).

Figure 4a shows the level of productive VDJ for each VH by cell type. Whilst mean levels are similar between μMT and normal pro-B cells, most VH in the latter are in either higher or lower productivity VDJ than the mean. This selection is strongly associated with clan and is maintained over the pre-B transition (Figure 4b).

**Figure 4.**
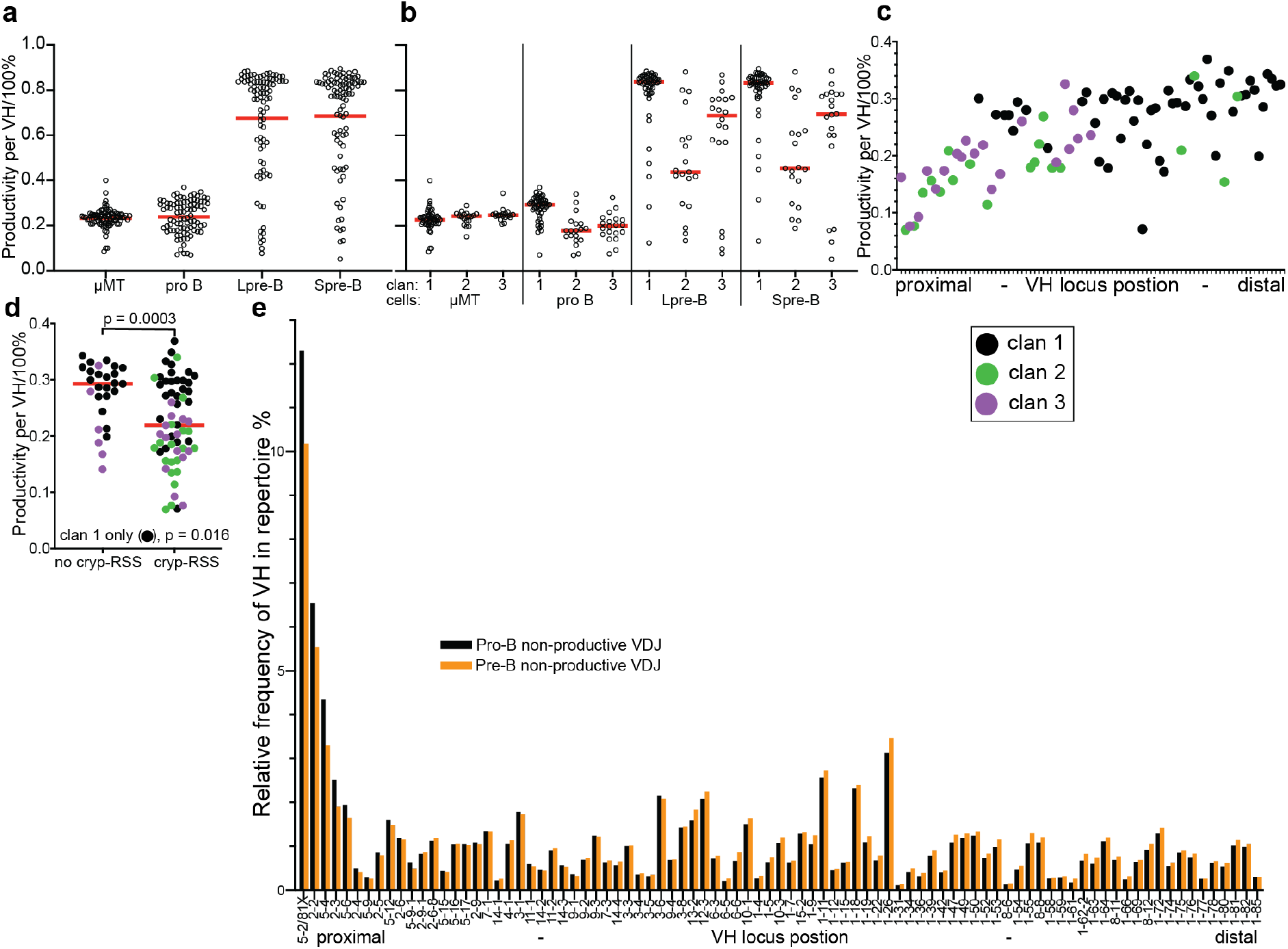
Selection and counter-selection of productive VDJ over the pre-B transition. **a**, VDJ productivity per VH for the four cell types analysed. mMT, mMT pro-B cells; pro **b**, pro-B cells, Lpre-B, large pre-B cells; Spre-B, small pre-B cells. Red bars show mean values. b, VDJ productivity per VH by VH clan, for the four cell types analysed. x-axis labels as previous panel. Red bars show median values. **c**, pro-B cell VDJ productivity per VH plotted by IgH locus position and coloured by clan – see key. **d**, VDJ productivity per VH for pro-B cell VDJ whose VH do and don’t have a cryptic-RSS. coloured by clan – see key. no cryp-RSS, no cryptic RSS; cryp-RSS, has cryptic RSS. Red bars show median values. Cryptic RSS defined as TACTGTG. **e**, Frequency of VH in non-productive VDJ from pro-B and small pre-B cells by locus position.

The association of VDJ productivity with clan is also a trend of VDJ productivity with VH locus position (Figure 4c).The lower productivity of proximal VH VDJs arguably could be because they recombined earlier and consequently the productive VDJ have exited the pro-B compartment before detection. However this is not the case, as it would result in equivalent productivity levels between clans in the pre-B compartments, which we do not observe (Figure 4b).

Importantly, most of the variation in pro-B cell VDJ productivity is absent in μMT pro-B cells (Figure 4a), implying that this VDJ productivity variation is largely driven by heavy-chain signalling, and not intrinsic variation between VH in forming productive VDJs.

### Productive VH5-2/81X rearrangements disappear from the pre-B cell repertoire

Currently, it is thought that if a first VDJ rearrangement is productive but cannot pair with the SLC, the second allele rearranges, and if this is functional then the cell will enter the pre-B transition with two productive heavy-chains. Non-pairing μ-chains are not toxic and such cells are thought to make up approximately 5% of post pre-B transition cells (Minegishi & Conley, 2001; ten Boekel et al., 1998). The data for VH5- 2/81X does not agree with this model. 8.8% of productive pro-B VDJ are VH5-2. As most of these do not pair, and the second allele is rearranged if available, these productive VH5-2 VDJ should enter the pre-B compartment as passengers. We only see 0.53% of the repertoire as productive VH5-2 VDJ in small pre-B cells, and this figure includes functional VH5-2/81X VDJ. It appears that many productive, non-pairing, VH5-2 VDJ are ‘disappearing’ from the repertoire.

We hypothesised, therefore, that productive, non-SLC-pairing, VH5-2 VDJ rearrangements, and likely others, become targets for VH-replacement and that this explains their disappearance.

### Low VDJ productivity correlates with possession of a cryptic RSS

Cryptic-RSSs facilitate VH-replacement, so we analysed their association with VDJ productivity. Low VDJ productivity correlates strongly with the presence of a cRSS in the VH (Figure 4d). This appears associated with clans 2 and 3, but clan 1 VH alone also show a significant correlation between cRSSs and low productivity (Fig. 4d). These results are consistent with VH replacement of productive, non-pairing, VDJ in pro-B cells, mediated by recombination at the cRSS.

### VH-Replacement frequency mirrors loss of productive VDJ

In contrast to normal VDJ-rearrangement excision-circles, where back-to-back RSSs are flanked by intergenic DNA, in VHR, cleavage at the cryptic RSS could produce an excision circle with the replaced VH adjacent to its cRSS, ligated to the RSS and 3’ intergenic sequence of the invading VH (Figure 5a). We designed a sequencing strategy to detect VHR circles, with the run-off primer position indicated on Figure 5a, ‘Seq.’

**Figure 5.**
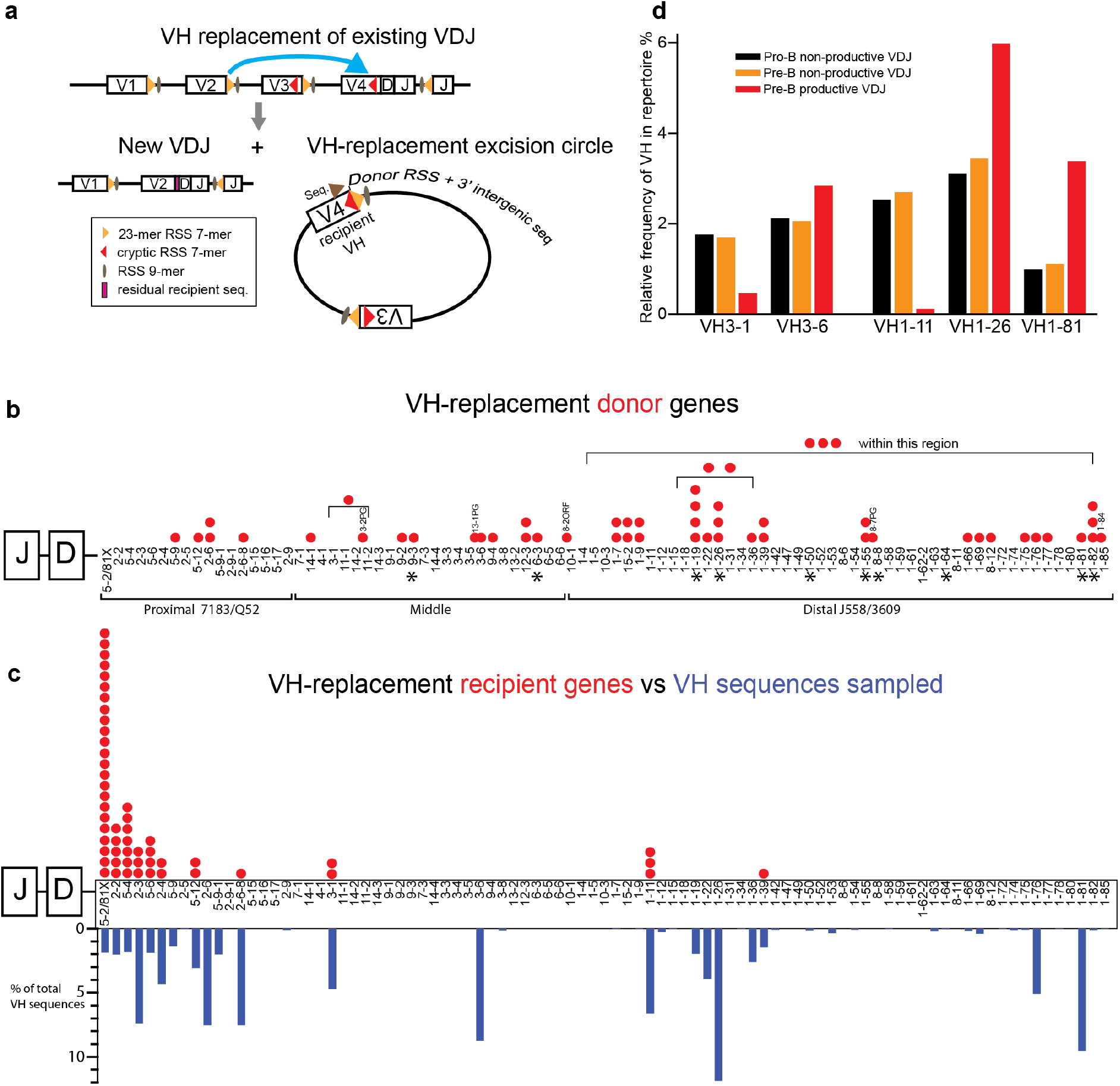
Location and frequency of use of VH-replacement donor and recipient VH. **a**, Diagram showing how VH replacement can generate an excision circle containing the replaced VH followed by the RSS and 3’ intergenic sequence of the invading/donor VH. ‘Seq.’ with brown arrow indicates approximate location of run-off primers for sequencing of excision circle to yield sequence that detects recipient and donor VH. **b**, Diagram of IgH locus indicating position and frequency of use of VH replacement donor VH genes, from the 53 VH replacement excision circles detected. Red dots indicate detection of a donor sequence from that VH. The brackets indicate the range of locations some of the donor VH may have, due to insufficient sequence data to precisely locate them. Asterisks indicate the ten VH that show the largest gains in productive VDJ over the pre- B transition. Some of the donor VH were open reading-frames (ORFs) or pseudogenes (PG), these are labelled above the dots indicating their locus position. In addition there is one VH donor, VH1-84, that is also indicated at the extreme distal end. It is not included in the main VH locus labelling because it is not one of the 89 most frequently used VH. **c**, Diagram of IgH locus indicating the position and frequency of use of VH replacement recipient VH genes, from the 53 VH replacement excision circles detected. Red dots indicate VH replacement events at that VH. The blue bars below the locus labelling indicate the VH sampling distribution of the total sequences performed. **d**, Frequency of selected VH in pro-B non-productive, pre-B non-productive and pre-B productive VDJ. Data extracted from Figure 2b and Figure 4e.

To avoid sequencing every VH from every cell, the sequencing primers focussed on VH more likely to have been replaced i.e. those that show a large drop in productive VDJ levels. These were VH5-2/4/6/12 and VH2-2/3/6/-6-8; and VH3-1 and VH1-11. VH1-11 shows the largest drop in productive VDJ for VH1s, which generally show strong positive selection (Figure 2b). As controls we included primers targeting the positively selected VH3-6, in contrast to VH3-1; and VH1-26 and 1-81.

We found 53 sequences in total in gDNA from 3 pro-B cell pools and a large pre-B cell pool. All sequences that fulfilled the above search criteria, for VH followed by back-to-back RSS (+/- insertions) followed by ectopic VH intergenic sequence, showed donor genes distal to recipients. The VH replacement circle sequences are in Supplementary document 1 and donor/recipient pairs are in Supplementary Table 1c. 18/53 sequences have insertions of between 1 and 4bp between the back-to-back 7mer RSSs.

The position of the donor/invading VH genes on the VH locus is shown in Figure 5b. The sequencing strategy is agnostic of the identity of donor genes (Fig5a). VH replacement donor VH are derived throughout the locus, but with a bias toward distal VH1. The ten VH showing the highest gain in productive VDJ (from Figure 2b) over the pre-B transition are marked with asterisks. We detect VHR donors from seven of these, associating VH replacement donors with high net gains in productive VDJ. Donor VH also include at least 3 pseudogenes and an open reading frame (ORF), VH8-2.

The position of the VH replacement recipient genes is shown in Figure 5c. The detection of these is non-exhaustive as we only probed for the candidates described above, with primers that sampled sequences in the proportions shown in blue bars. VH5-2/81X, which shows the highest loss of productive VDJ, almost all of which cannot pair with the SLC (ten Boekel et al. 1997), dominates as a target for VH replacement. The next four proximal VH, which also show high losses of productive VDJ, are also frequently replaced, compared to the other proximal VH sequenced. Importantly, for VH3-1 and VH1-11, which show large losses of productive VDJ, but no losses of non-productive VDJ (Figure 5d), we detected 2 and 3 replacement events, but none for the controls

VH3-6 and VH1-26/81, which gain productive VDJ at the pre-B transition. We conclude that VHR drives the removal of productive non-pairing VDJ from the pro-B cell repertoire.

## Discussion

Tying VH-Replacement to removal of non-pairing VDJ rearrangements defines a physiological role for VHR, and resolves a long-lasting question fundamental to VDJ repertoire formation, that of what happens to the non-pairing VDJ (Marshall et al., 1996; ten Boekel et al., 1997). This also explains at least some of the distal repertoire shift, as we show that distal VH are more tolerant of CDR3 composition, thus can replace productive non-pairing proximal VDJs.

While our strategy cannot determine whether the replaced VH was productive or non-productive, VH3-1 and VH1-11, for which we detect VHR, only demonstrate loss of productive VDJ over the pre-B transition (Figure 5d). With VH5-2/81X, the loss of productive VDJ (Figure 2b), which hardly ever pair, is six-fold higher than non-productive VDJ (Figure 4e). This is consistent with VH-replacement with a strong bias toward productive VDJ.

Whilst productive non-pairing VDJ may be no better molecular substrates for VHR, they may be more available for secondary recombination. Non-paired μ-chains can still signal allelic exclusion (Shimizu et al., 2002), whilst at the same time incompletely supressing Rag and TdT (Galler et al., 2004). This would favour VH replacement over second allele recombination; as compared to a non-productive VDJ - which may stop continued recombination on the same allele through nonsense-mediated decay signalling (Fuxa et al., 2004; Roldan et al., 2005). μMT mice do not undergo the VDJ selection (Figure 4a) that we show is impacted by VHR, confirming that this selection is driven by μ-chain signalling and thus productive VDJ expression.

μ-chains vary in their dependency on SLC properties for effective signalling (Kohler et al., 2008). Productive non-pairing μ-chains may well also vary in their residual signalling capacity. Some can signal autonomously (Galler et al., 2004) and may support developmental progress, such as the auto-reactive heavy-chains found in SLC −/− mice (Keenan et al., 2008). Some heavy-chain only antibodies can even appear in the periphery (Zou et al., 2007). However, most unpaired μ-chains will signal less than a functional pre-BCR at least temporarily stalling the cell which could then be rescued by VHR.

Our detection of VHR greatly underestimates the number of events. Firstly, most VHR will result in non-productive VDJ. This will lead to cell death or second allele rearrangement with similar chance of failure. Secondly, our detection of VHR is dependent on detection of ligated excision circles. Even the fate of the un-ligated signal joint ends in normal V-DJ recombination is unclear (Lieber, 2010). Further, up to 20% of peripheral B-cell VDJ can be derived from VHR in a mouse with a monoclonal, functional VDJ, despite a shortened pro-B cell transit (Kumar et al., 2015; Sun et al., 2015). This suggests that VHR can account for all the changes in productive VDJ we observe, considering the likely extended pro-B cell phase of a non-pairing productive VDJ in a normal physiological context.

That we detect VH-Replacement donor VH from right across the *IgH* locus, in a physiological context, is also consistent with a developmental stalling of B-cells with productive non-pairing VDJ. Such stalling would allow the locus contraction necessary to facilitate distal VH to be donors for VH-Replacement of a more proximal VDJ. In contrast, a mouse knock-in model using a functional VDJ showed only proximal VH as donors for VH-Replacement, consistent with an accelerated pro-B cell transit (Sun et al., 2015).

We have shown independent selection of three CDR3 sequence features over the pre-B transition: N-terminal amino acids, RF, and strong VH-specific counter-selection of JH. The selection of the VH encoded N-terminal A97 and R98 is VH-universal, highlighting the importance of these residues for successful pre-B transition, as is the counter- selection of RF2. Conversely, the strong VH-specific counter-selection we observe, that removes a third of the pro-B cell VDJ repertoire, is associated either with JH2/3 in VH2/3/5 or with specific CDR3 residues, e.g. G97 in VH1-11 or S/K98 in VH2s. We suggest these features impact SLC-pairing the most, driving most of the VH- replacement, excepting for VH5-2/81X, which largely fails to pair.

The bone-marrow cytokine milieu is altered by ageing and chronic inflammation (Dowery et al., 2021; Koohy et al., 2018; Pioli et al., 2019). Inflammatory stimuli can drive release of B-cell progenitors into the periphery (Nagaoka et al., 2000; Ueda et al., 2004). These stresses significantly alter the pre-B transition, which has impacts on the peripheral repertoire. Our in-depth study of the structure and selection of the normal heavy-chain repertoire provides a good foundation to further investigate such alterations.

## Methods

### Mice

Wild-Type C57BL/6Babr and μMT x B6SJL mice were maintained in the Babraham Institute Biological Services Unit in accordance with the institute Animal Welfare and Ethical Review Body and Home Office Rules under Project License 80/2529. ARRIVE guidelines were followed. All mice were males between 11.7 and 14 weeks old.

### Primary Cells

Mice were euthanised with CO2 asphyxiation followed by cervical dislocation, or cervical dislocation followed by pithing. Wild-type bone marrow was flushed from femurs and tibias, washed once with PBS and subject to non-B cell depletion using biotinylated antibodies against CD11b (MAC-1; ebioscience; 1:1600), Ly6G (Gr-1; ebioscience; 1:1600), Ter119 (ebioscience; 1:400) and CD3e (ebioscience; 1:800), followed by incubation on ice for 30 minutes. Cells were washed and Streptavidin MACS beads were then added (5 μl/10^7^ cells in 100 μl; Miltenyi) and incubated at 4deg for 15 minutes with occasional mixing. MACS LS columns were equilibrated and washed cells were loaded with the flow through collected for flow sorting. After flushing

μMT mouse bone marrow cells were washed once with PBS and subject to positive selection for CD19 using MACS LS columns (Miltenyi) according to manufacturers instructions. Cells were then used directly for genomic DNA extraction

### FACS

Mouse bone-marrow (BM) B-cell progenitors were sorted from femurs and tibias, for at least two biological replicate pools of 4 or more, 12–14-week-old, C57BL/6J male mice (Figure S1a). B-cell enriched wild-type bone marrow cells were Fc Blocked for 10 minutes (1:400, eBioscience, clone 93) then stained for flow sorting with the following markers: BV421 anti-CD19 (1:400, BD Biosciences, clone 1D3), BV650 anti-CD93 (1:800, BD Biosciences, cloneAA4.1), PE-Cy5 anti-CD25 (1:500, eBioscience, clonePC61.5), ef660 anti-IgM (1:400, eBioscience, clone11/41), PE anti-IL7Ra (1:400, eBioscience, clone eBioSB/199), PE-Cy7 anti-CD43 (1:100, BD Biosciences, cloneS7), FITC anti-CD24 (1:800, eBioscience, clone 30F1), APCeFluor780 viability dye (1:2000, eBioscience). When staining for cytoplasmic μ-chain was also done, cells were stained according to the above panel but using BUV737 anti-CD19 (1:400, BD Biosciences, clone 1D3). Cells were washed, fixed using Cytofix/Cytoperm (BD Biosciences) and stained for 30 minutes on ice with ef450 anti-IgM (1:400, Invitrogen, clone eb121-15F9). Cells were sorted on a BD FACSAria Fusion or FACSAria II. Flow sorter gating strategy is shown in Figure S1a. The gating strategy combines and extends previous strategies. The phenotypes were: All cells, CD19+, CD93+; pro-B cells (Hardy fraction BC, Basel pre-BI), CD25-, IgM-, IL-7R+, CD43hi, CD24mid; large pre-B cells (Hardy fraction C’, Basel large pre-BII) CD25-, IgM-, IL-7R+ CD43mid/lo, CD24hi; small pre-B cells (Hardy fraction D, Basel small pre-BII), CD25+, IgM-. We have analysed c-kit+ (clone 2B8) CD43+, IgM- pro-B cells and find some c-kit+ cells have a large pre-B cell phenotype (Figure S1b). Pro-B subset gates, using cytoplasmic m-chain staining, are shown in Figure 3c. Cells were then centrifuged at 450g for 5 minutes, resuspended in 1 x DNA/RNA shield and frozen at −20deg.

### VDJseq

Mouse bone-marrow (BM) B-cell progenitors were sorted from femurs and tibias, for at least two biological replicate pools of 4 or more, 12–14-week-old, C57BL/6J male or μMT x B6SJL male mice.

Genomic DNA from individual progenitor cell pools was subject to quantitative VDJ repertoire analysis (Chovanec et al., 2018). This detects both DJ and VDJ recombinants allowing measurement of the frequency of VDJs in populations.

Libraries were sequenced on an Illumina MiSeq. Sequence reads were analysed using the Babraham Linkon pipeline (https://github.com/peterch405/BabrahamLinkON).

After processing sequence data using the BabrahamLinkon pipeline, fasta files from the penultimate step were also processed through the IMGT/HighV-QUEST web service to extract D-region reading frames (RFs) which were then merged with main data using sequence ID in R. We use the D-reading frame convention of Ichihara et al. (Ichihara et al., 1989), not IMGT, so RF i.d’s were transposed 1 to 2, 2 to 3, 3 to 1.

Of the 195 C57BL/6 VH genes, 123 are recombinationally active (Bolland et al., 2016). We focused on VH recombinants detected at >0.1% frequency, to allow reproducible comparisons from datasets of varying sizes. These 89 VH (supplementary table 1b) represent approximately 98% of productive VDJ recombinants in a pro B-cell repertoire.

To address a few minor discrepancies in the IMGT VH gene reference database, we have used a slightly modified VH reference database in the BabrahamLinkon pipeline used to analyse VDJseq data, which can be called using custom_ref during clone assembly. https://github.com/peterch405/BabrahamLinkON/tree/master/babrahamlinkon/resources/IgBlast_database We observed strong correlation of VH frequency between biological replicates (R^2^ =0.98-0.99), (Figure S1c), so replicate sequence pools were merged.

All downstream analysis was performed using RStudio to generate spreadsheets and/or data suitable for using in Graphpad Prism, which was used for plotting and statistical analysis. Significance testing was performed with an un-paired two-tailed t-test with Welch’s correction and correlation was measured with simple linear regression.

### VH replacement excision circle sequencing

Excision circles were sequenced using an adapted VDJseq protocol. We used nested VH framework-3 primer pools for the single cycle run-offs, followed by the first PCR step of the VDJseq protocol. Sequences shown in Supplementary table 1d. These targeted VH2-2,2-3,2-6,2-6-8,5-2,5-4,5-6,5-12,1-11,1-26,1-81,3-1,3-6. The run-off annealing temperature was changed to 56deg., and the PCR1 annealing temperature to 60deg, with 12 or 13 cycles depending on starting DNA amounts. Library preparation and sequencing then resumed as for VDJseq. Libraries were sequenced on an Illumina MiSeq or an Element Biosciences Aviti.

Most sequences originate from the germline VH genes present in every cell. If VH replacement occurs and resolves in a manner related to normal VDJ recombination, there should be VH-replacement circles, with VH sequence followed by a head-to-head 7-mer followed by ectopic intergenic sequence 3’ from the invading VH.

Raw sequences were searched for VH replacement circles in two ways. The main approach was to assemble the paired end reads using PEAR, reverse and complement them, and deduplicate using the UMIs and sequence length. We then ran sequences through a bespoke script to detect replacement circles. This searches for 60bp of framework-3 sequence of any particular VH followed by a variable gap that includes the back to back 7-mer RSSs, which allows for imprecise joining, and then for 60bp of the intergenic sequence 3’ of any VH that is not the same as the framework-3 VH. These scripts are available here: https://github.com/s-andrews/replacementcircles Sequences were then manually curated to confirm the presence of the back-to-back cryptic 7-mer/normal 7-mer RSSs and confirm the identity of the VH sequences involved using IgBlast. As an adjunct, to search reads that didn’t extend to the end of the second VH search region, we also searched deduplicated reads directly for back-to-back cryptic 7-mer/normal 7-mer RSS sequences, using grep in shell script, allowing for up to 3bp insertions between RSSs. The most common back-to-back cryptic 7- mer/normal 7-mer RSS sequence is TACTGTG/CACAGTG, although there is variation in the normal VH RSS 7-mer necessitating a redundancy in this search, TACTGTG/CACA[GAT]TG. Returned sequences not present from the main search were manually curated in the same way.

## Acknowledgements

The authors thank all the members of the Babraham Institute Flow Cytometry facility, Biological Services Unit, Bioinformatics Group and Genomics Facility. We thank Alex Whale and Jon Houseley for valuable discussion. We also thank Martin Turner for reviewing the manuscript.

## Competing Interests

The authors declare no competing interests

## Materials and Correspondence

Correspondence and material requests should be directed toward Harry N White.

## Author Contributions

HNW Conceptualisation, Validation, Investigation, Data Curation, Formal Analysis, Visualisation, Software, Methodology, Project Administration, Writing-original draft, Writing-review and editing

PC Methodology, Software, Writing-review and editing

LB Software, Data Curation, Formal Analysis, Visualisation

ECF Investigation

GB Investigation

SA Software, Data curation, Formal Analysis, Conceptualisation

AEC Conceptualisation, Validation, Funding Acquisition, Project Administration, Supervision, Writing-review and editing

## Data Availability

The VDJ-seq and VH-replacement circle raw sequencing files generated in this study have been deposited in SRA under Bioproject ID PRJNA1194793 and are publicly available.

## Notes

### Competing Interest Statement

The authors have declared no competing interest.

