## Supplemental data for "VH-replacement shapes the antibody repertoire by targeting non-pairing heavy-chains"

### Figure S1

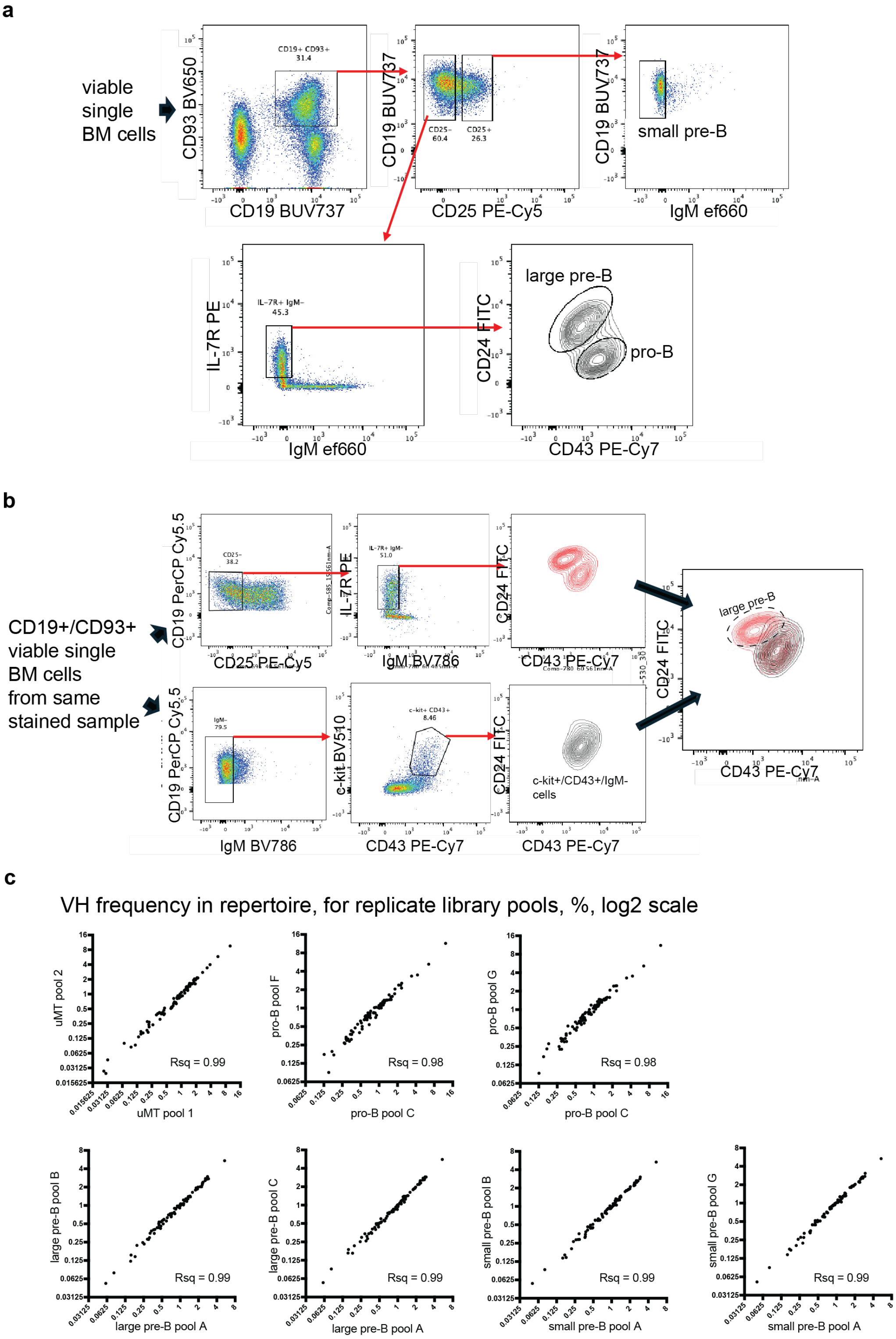

**Figure S1. FACS gating strategy and VDJseq library replicate comparisons**
**a**, Babraham Institute FACS gating strategy for selection of pro-B cells, large pre-B cells and small pre-B cells. **b**, Diagram showing that some c-kit+/CD43+/IgM- bone-marrow B-cells overlap with CD25-/IL-7R+/IgM-/CD24+/CD43lo gate for large pre-B cells. **c**, Frequency of individual VH in VDJseq libraries from biological replicate libraries from other mouse bone marrow pools. Consult Supplementary Table 1a for composition of bone marrow B-cell pools

### Figure S2

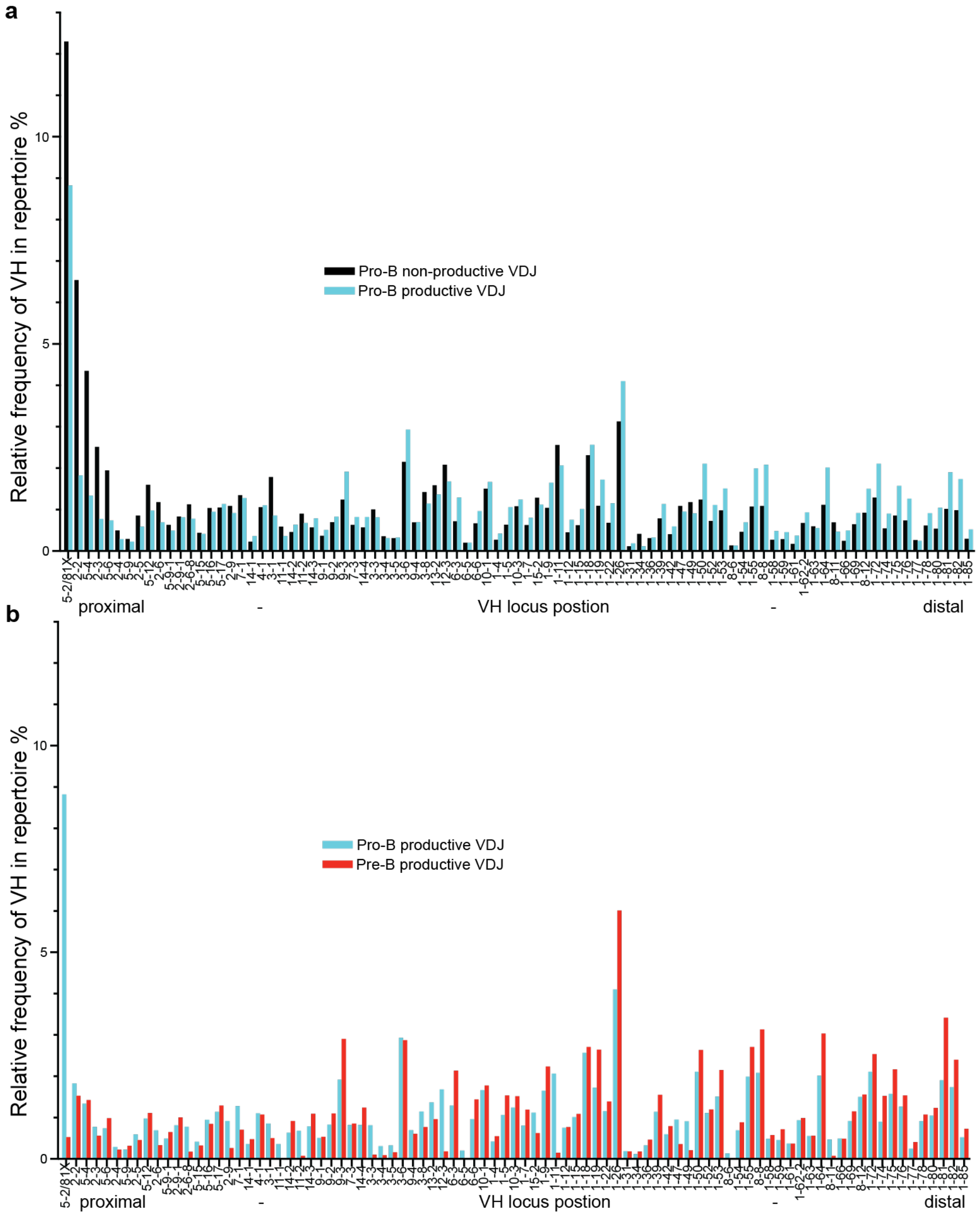

**Figure S2. The two measured steps of VH selection over the pre-B transition**

The overall change in VH frequencies over the pre-B transition is shown in Figure 2b. Through VDJ seq this can be broken down into two steps, the change between non-productive and productive VH/VDJ levels in pro-B cells, and the change in productive VH/VDJ levels between pro- and pre-B cells. **a**, VH frequency in non-productive and productive VDJ from pro-B cells, by locus position. **b**, VH frequency in productive VDJ from pro-B cells and pre-B cells, by locus position. **c**, Change in VH frequency between non-productive and productive VDJ from pro-B cells versus change in frequency between productive VDJ from pro- and pre-B cells, coloured by clan. R-squared values for the correlation are show with and without the five most proximal VH.

### Figure S3

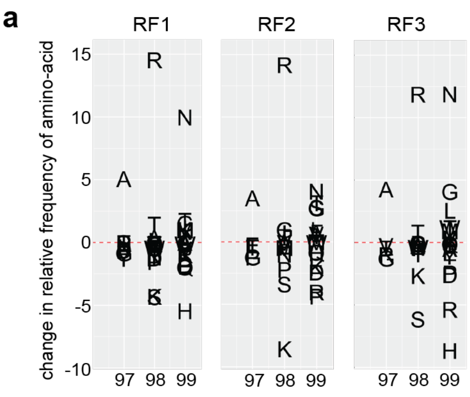

**b** Arginines per CDR3

| | pro-B | small pre-B |
|---|---|---|
| **VH1** | 1.11 | 1.16 |
| **VH2** | 0.83 | 0.9 |
| **VH5** | 1.31 | 1.32 |
| n | 19518 | 259397 |

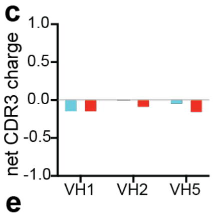

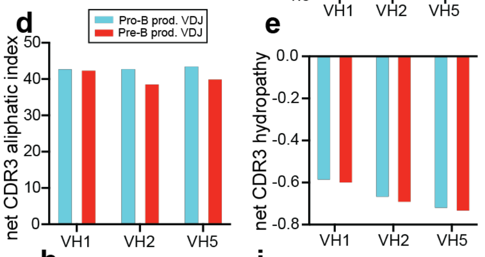

**f** Argines per RF1 CDR3

| | proB/if/stop | pro-B | small pre-B |
|---|---|---|---|
| **VH1** | 1.01 | 1.03 | 1.07 |
| **VH2** | 0.76 | 0.71 | 0.82 |
| **VH5** | 1.05 | 1.18 | 1.222 |
| n | 1259 | 8444 | 130932 |

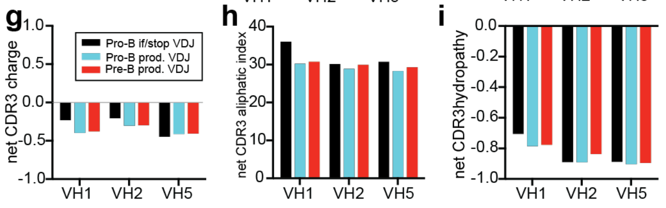

**Figure S3. Consistent selection of N-terminal CDR3 amino-acids in all D reading frames, and biophysical properties of CDR3s, over the pre-B transition**

**a**, Change in frequency of the first three CDR3 amino-acid residues over the pre-B transition by D-reading-frame. Data shown here for VH2 only which also shows the strongest Y101 selection and RF2 counter-selection. **b**, Mean number of Arginine residues per CDR3 for VH1, VH2, VH5 families from pro-B productive VDJ and pre-B productive VDJ. **c**, net CDR3 charge for VH1, VH2, VH5 families from pro-B productive VDJ and pre-B productive VDJ. **d**, net CDR3 aliphatic index for VH1, VH2, VH5 families from pro-B productive VDJ and pre-B productive VDJ. **e**, net CDR3 hydropathy for VH1, VH2, VH5 families from pro-B productive VDJ and pre-B productive VDJ. **f**, Mean number of Arginine residues per RF1 CDR3 for VH1, VH2, VH5 families from pro-B non-productive (in-frame but with a stop codon), pro-B productive VDJ and pre-B productive VDJ. **g**, net RF1 CDR3 charge for VH1, VH2, VH5 families from pro-B non-productive (in-frame but with a stop codon), pro-B productive VDJ and pre-B productive VDJ. **h**, net RF1 CDR3 aliphatic index for VH1, VH2, VH5 families from pro-B non-productive (in-frame but with a stop codon), pro-B productive VDJ and pre-B productive VDJ. **i**, net RF1 CDR3 hydropathy for VH1, VH2, VH5 families from pro-B non-productive (in-frame but with a stop codon), pro-B productive VDJ and pre-B productive VDJ. Values for c, d, e, g, h, i were calculated using the method and algorithm from Protparam, https://web.expasy.org/protparam.

### Supplementary Table 1

# a

### VDJseq library metadata

| Cells | | Mouse BM pool | Mice/pool | VDJ reads | DJ reads | VDJ:DJ | proportion alleles VDJ | VDJ alleles/cell | Prod. Seqs | VDJ Productive % |
|---|---|---|---|---|---|---|---|---|---|---|
| **muMT proB** | | 1 | 4 | 45138 | 62081 | 0.73 | 0.420988817 | 0.841977635 | 10480 | 0.23217688 |
| | | 2 | 4 | 44102 | 59923 | 0.74 | 0.42395578 | 0.84791156 | 10185 | 0.230941907 |
| | mean | | | | | | | | | 0.231559394 |
| **ProB** | | C | 8 | 28267 | 47208 | 0.6 | 0.374521365 | 0.749042729 | 5755 | 0.203594297 |
| | | F | 6 | 37777 | 56317 | 0.67 | 0.401481497 | 0.802962994 | 8124 | 0.215051486 |
| | | G | 6 | 36854 | 53360 | 0.69 | 0.408517525 | 0.81703505 | 7330 | 0.198892929 |
| | mean | | | | | | | | | 0.205846237 |
| **Large PreB** | | A | 4 | 142139 | 41803 | 3.4 | 0.772738146 | 1.545476291 | 102352 | 0.720083862 |
| | | B | 4 | 69917 | 21269 | 3.29 | 0.766751475 | 1.53350295 | 50135 | 0.717064519 |
| | | C | 8 | 107894 | 29338 | 3.68 | 0.786216043 | 1.572432086 | 77423 | 0.717583925 |
| | mean | | | | | | | | | 0.718244102 |
| **Small PreB** | | A | 4 | 125989 | 36802 | 3.42 | 0.773930991 | 1.547861983 | 91392 | 0.725396662 |
| | | B | 4 | 107739 | 32224 | 3.34 | 0.769767724 | 1.539535449 | 77684 | 0.721038807 |
| | | G | 6 | 135015 | 38950 | 3.47 | 0.776104389 | 1.552208778 | 96572 | 0.715268674 |
| | mean | | | | | | | | | 0.720568047 |

| Pro-B subsets | | Mice/pool | VDJ reads | DJ reads | VDJ:DJ | proportion alleles VDJ | VDJ alleles/cell | Prod. Seqs | VDJ Productive % |
|---|---|---|---|---|---|---|---|---|---|
| | Large cyto-mu neg. | 20 | 12480 | 76492 | 0.16315432 | 0.140268849 | 0.280537697 | 708 | 0.056730769 |
| | Small cyto-mu neg. | 20 | 51873 | 52119 | 0.99528003 | 0.498817217 | 0.997634433 | 3794 | 0.073140169 |
| | Small cyto-mu pos. | 20 | 32740 | 15549 | 2.10560165 | 0.678001201 | 1.356002402 | 22615 | 0.690745266 |
| | Large cyto-mu pos. | 20 | 17241 | 12904 | 1.33609733 | 0.571935644 | 1.143871289 | 12388 | 0.718519807 |

**b**

### 89 reliably detectable VH

| VCALL | Classic name | Clan |
|---|---|---|
| IGHV5-2 | 7183.2.3_(81X) | 3 |
| IGHV2-2 | Q52.2.4 | 2 |
| IGHV5-4 | 7183.4.6 | 3 |
| IGHV2-3 | Q52.3.8 | 2 |
| IGHV5-6 | 7183.7.10 | 3 |
| IGHV2-4 | Q52.5.13 | 2 |
| IGHV5-9 | 7183.9.15 | 3 |
| IGHV2-5 | Q52.7.18 | 2 |
| IGHV5-12 | 7183.12.20 | 3 |
| IGHV2-6 | Q52.8.22 | 2 |
| IGHV5-9-1 | 7183.14.25 | 3 |
| IGHV2-9-1 | Q52.9.29 | 2 |
| IGHV2-6-8 | Q52.10.33 | 2 |
| IGHV5-15 | 7183.18.35 | 3 |
| IGHV5-16 | 7183.19.36 | 3 |
| IGHV5-17 | 7183.20.37 | 3 |
| IGHV2-9 | Q52.13.40 | 2 |
| IGHV7-1 | S107.1.42 | 3 |
| IGHV14-1 | SM7.1.44 | 1 |
| IGHV4-1 | X24.1.45 | 3 |
| IGHV3-1 | 36-60.1.46 | 2 |
| IGHV11-1 | VH11.1.48 | 3 |
| IGHV14-2 | SM7.2.49 | 1 |
| IGHV11-2 | VH11.2.53 | 3 |
| IGHV14-3 | SM7.3.54 | 1 |
| IGHV9-1 | VGAM3.8-1-57 | 1 |
| IGHV9-2 | VGAM3.8-2-59 | 1 |
| IGHV9-3 | VGAM3.8-3-61 | 1 |
| IGHV7-3 | S107.3.62 | 3 |
| IGHV14-4 | SM7.4.63 | 1 |
| IGHV3-3 | 36-60.3.64 | 2 |
| IGHV3-4 | 36-60.4.66 | 2 |
| IGHV3-5 | 36-60.5.67 | 2 |
| IGHV3-6 | 36-60.6.70 | 2 |
| IGHV9-4 | VGAM3.8-4-71 | 1 |
| IGHV3-8 | 36-60.8.74 | 2 |
| IGHV13-2 | 3609N.2.77 | 3 |
| IGHV12-3 | VH12.1.78 | 2 |
| IGHV6-3 | J606.1.79 | 3 |
| IGHV6-5 | J606.3.81 | 3 |
| IGHV6-6 | J606.4.82 | 3 |
| IGHV10-1 | VH10.1.86 | 3 |
| IGHV1-4 | J558.2.88 | 1 |
| IGHV1-5 | J558.3.90 | 1 |
| IGHV10-3 | VH10.3.91 | 3 |
| IGHV1-7 | J558.4.93 | 1 |
| IGHV15-2 | VH15.1.95 | 1 |
| IGHV1-9 | J558.6.96 | 1 |
| IGHV1-11 | J558.8.98 | 1 |
| IGHV1-12 | J558.9.99 | 1 |
| IGHV1-15 | J558.12.102 | 1 |
| IGHV1-18 | J558.16.106 | 1 |
| IGHV1-19 | J558.18.108 | 1 |
| IGHV1-22 | J558.22.112 | 1 |
| IGHV1-26 | J558.26.116 | 1 |
| IGHV1-31 | J558.31.121 | 1 |
| IGHV1-34 | J558.34.124 | 1 |
| IGHV1-36 | J558.36.126 | 1 |
| IGHV1-39 | J558.39.129 | 1 |
| IGHV1-42 | J558.42.132 | 1 |
| IGHV1-47 | J558.47.137 | 1 |
| IGHV1-49 | J558.49.141 | 1 |
| IGHV1-50 | J558.50.143 | 1 |
| IGHV1-52 | J558.52.145 | 1 |
| IGHV1-53 | J558.53.146 | 1 |
| IGHV8-6 | 3609.5.147 | 2 |
| IGHV1-54 | J558.54.148 | 1 |
| IGHV1-55 | J558.55.149 | 1 |
| IGHV8-8 | 3609.7.153 | 2 |
| IGHV1-58 | J558.58.154 | 1 |
| IGHV1-59 | J558.59.155 | 1 |
| IGHV1-61 | J558.61.157 | 1 |
| IGHV1-62-2 | J558.64.162 | 1 |
| IGHV1-63 | J558.66.165 | 1 |
| IGHV1-64 | J558.67.166 | 1 |
| IGHV8-11 | 3609.11.169 | 2 |
| IGHV1-66 | J558.69.170 | 1 |
| IGHV1-69 | J558.72.173 | 1 |
| IGHV8-12 | 3609.12.174 | 2 |
| IGHV1-72 | J558.75.177 | 1 |
| IGHV1-74 | J558.77.180 | 1 |
| IGHV1-75 | J558.78.182 | 1 |
| IGHV1-76 | J558.79.184 | 1 |
| IGHV1-77 | J558.80.186 | 1 |
| IGHV1-78 | J558.81.187 | 1 |
| IGHV1-80 | J558.83.189 | 1 |
| IGHV1-81 | J558.84.190 | 1 |
| IGHV1-82 | J558.85.191 | 1 |
| IGHV1-85 | J558.88.194 | 1 |

c

### VH-replacement donors and recipients

| Sample | Recipient | Donor | Insertions |
|---|---|---|---|
| LP2 | 5-2 | 1-55 | 2 |
| SN2 | 5-4 | 2-6-8 | 0 |
| SN2 | 5-6 | 15-2 | 0 |
| SN2 | 5-2 | 3-2 PG | 0 |
| SN2 | 5-2 | 1-66 | 0 |
| SN2 | 5-2 | 1-x | 1 |
| SN2 | 5-2 | 1-x | 1 |
| SN2 | 5-4 | 3-1/2PG | 2 |
| SN2 | 5-12 | 2-6 | 3 |
| SN2 | 5-6 | 5-9 | 3 |
| SP2 | 5-2 | 1-19 | 0 |
| SP2 | 5-4 | 1-19/36 | 0 |
| DEBC | 5-2 | 1-39 | 4 |
| DEBC | 5-2 | 12-3 | 0 |
| DEBC | 2-6-8 | 1-26 | 1 |
| DEBC | 2-3 | 1-7 | 0 |
| DEBC | 5-2 | 1-76 | 0 |
| DEBC | 5-6 | 1-81 | 0 |
| DEBC | 2-4 | 1-82 | 3 |
| DEBC | 5-2 | 1-36 | 0 |
| DEBC | 5-2 | 3-6 | 0 |
| DEBC | 5-4 | 13-1 PG | 0 |
| DEBC | 1-11 | 1-84 | 0 |
| LP3 | 2-3 | 1-82 | 0 |
| LP3 | 1-11 | 1-19 | 0 |
| LP3 | 5-4 | 1-x | 3 |
| SN3 | 5-6 | 9-4 | 2 |
| SN3 | 1-39 | 1-75 | 2 |
| SN3 | 1-11 | 8-12 | 4 |
| SN3 | 5-2 | 1-77 | 0 |
| SN3 | 5-4 | 1-26 | 0 |
| SN3 | 5-2 | 1-82 | 0 |
| SN3 | 3-1 | 1-69 | 4 |
| SN3 | 2-2 | 1-7 | 0 |
| SN3 | 2-4 | 5-12 | 0 |
| SP3 | 5-2 | 1-55 | 0 |
| SP3 | 3-1 | 15-2 | 0 |
| SP3 | 5-2 | 1-9 | 0 |
| SP3 | 5-2 | 12-3 | 0 |
| SP3 | 2-3 | 1-9 | 0 |
| SP3 | 5-2 | 6-3 | 3 |
| SP3 | 5-2 | 8-7 PG | 0 |
| Cp | 5-4 | 9-2 | 2 |
| Cp | 2-2 | 1-26 | 0 |
| Cp | 2-2 | 1-19 | 0 |
| Cp | 2-2 | 1-22 | 0 |
| Cp | 5-2 | 1-39 | 0 |
| Cp | 5-2 | 14-1 | 0 |
| Cp | 2-2 | 9-3 | 4 |
| Cp | 5-2 | 1-19 | 3 |
| Cp | 5-2 | 8-2 ORF | 0 |
| Cp | 5-12 | 2-6 | 0 |
| Cp | 5-2 | 1-18to36 | 0 |

**d**

### primers for VH replacement circle sequencing

| Name | Sequence |
|---|---|
| VH1-11bio | CCAGTAAGTGGTGAAACTAACTACAATC |
| VH1-11P5 | GTGACTGGAGTTCAGACGTGTGCTCTTCCGATCTCATGGGCAAGGCCACATTCTCT*G |
| VH2-2bio | GGAGTGGTGGAAGCACAGAC |
| VH2-3bio | GGGGTGACGGGAGCACAAAT |
| VH2-61bio | TAATATGGGGTGTTGGAAGCACAAAT |
| VH2-62bio | ATATGGAGTGATGGAAGCACAACC |
| VH2-2/3/6 | GTGACTGGAGTTCAGACGTGTGCTCTTCCGATCTTCAWATCCAGACTGAGCATCAGCAA*G |
| VH5-2bio | GATGGTGGTAGCACCTACTATCC |
| VH5-4/6bi | TGGTAGTTACACCTACTATCCAGAC |
| VH5-12bio | GTGGTAGCACCTATTATCCAGAC |
| VH5-2P5 | GTGACTGGAGTTCAGACGTGTGCTCTTCCGATCTGAGAGACGATTCATCATCTCCAGA*G |
| VH5-4/6/1 | GTGACTGGAGTTCAGACGTGTGCTCTTCCGATCTGGSCGATTCACCATCTCCAGA*G |
| VH3-1/6bi | TGGTAGCAMTAACTACAACCCATC |
| VH3-1/6P5 | GTGACTGGAGTTCAGACGTGTGCTCTTCCGATCTCYCTCAAAARTCGAATCTCCATCACT*C |
| VH1-26bio | ATTAATCCTAACAATGGTGGTACTAGC |
| VH1-81bio | CCTAGAAGTGGTAATACTTACTACAATG |
| VH1-26/81 | GTGACTGGAGTTCAGACGTGTGCTCTTCCGATCTAGTTCAAGGGCAAGGCCACAYT*G |

**Supplementary Document 1: VH-replacement circle sequences**

**Key: back to back heptamers +/- insertion, donor/invader nonamer**

**Sort 2: RCseq after VDJseq, MiSeq Sequencing, shell-based searches for back to back RSS with up to 3 insertions. Pro-B subsets.**

**LP**

>M02293:67:000000000-LFYL9:1:1113:11780:16646 1:N:0:0 GTCCGC reverse complement 5-2 by 1-55 2 insertions
GGGCGATTCACCATCTCCAGAGACAATACCAAGAAGACCCTGTACCTGCAAATGAGCAGT
CTGAGGTCTGAGGACACAGCCTTGTATTACTGTGCCCACAGTGTTGCAACCACATCCTGA
GAGTGTCCAAAACCCTGGAGGAGTAGCAAACTGCCCTGGGACT

**SN**

>M02293:67:000000000-LFYL9:1:1112:14893:17472 1:N:0:0 ATGTCA reverse complement 5-4 by 2-6-8 (by blast) 0 insertions
GAGAGCGATTCTCATCTCCAGAGACAATGCCAAGAACAACCTGTACCTGCAAATGAGCCA
TCTGAAGTCTGAGGACACAGCCATGTATTACTGTGCACAGTGAGGGAAGTCCAATGTGAG
CCTGCACAAATACTTCTCTGCAGGGATGATCACAACCAGCAGGGGGCGCTGAGGATCCAA
AGGGACT

>M02293:67:000000000-LFYL9:1:1113:22579:13743 1:N:0:0 ATGTCA reverse complement 5-6 by 15-2 0 insertions
GGGCGATTCACCATCTCCAGAGACAATGCCAAGAACACCCTGTACCTGCAAATGAGCAGT
CTGAAGTCTGAGGACACAGCCATGTATTACTGTGCACAGTGTAACAGCTCATATCTGAAG
CATGTCAAAAAGTCTGAAGGCAGGAAGCTATTTTGGGTCTGATATTACCACAAAGATTAA
CCAAAGCAGTTGCTCAAAGCCTGTGGTTAAATGTTGAAGGAAAGA

>M02293:67:000000000-LFYL9:1:1119:25597:9060 1:N:0:0 ATGTCA reverse complement 5-2 by 3-2 Pseudogene 0 insertions
GGGCGATTCACCATCTCCAGAGACAATACCAAGAAGACCCTGTACCTGCAAATGAGCAGT
CTGAGGTCTGAGGACACAGCCTTGTATTACTGTGCACAGTGTGGAGTCTTCACTGTGAGC
CCAGACAAAAACCTCCTTGCAGAGCAGCACTGCACCAACAGGGGGCATGGAGCATACACC
AATAATGGGAAATCTACTTTAAAGTAGGTAAAAAAAAATTGTTGCCCAAAACTCTGGCAC
CAGACTCCAGGTATACAGTAAGAAACATTGGTGCTAATAATTTCAGTCAATATAATGTGC
ACAGCCCAATAAAACTCCTTATGTTATACCCAGAATTCTTCAATGGGAGCATCTTCCAGA
GCTTCCAAATCCCAGCTCAAAACCCAGCTGAAGTCTGACGGCCTCATAGAGGACAAATCA
TT

>M02293:67:000000000-LFYL9:1:2104:3918:15300 1:N:0:0 ATGTCA reverse complement 5-2 by 1-66 0 insertions
GAGAGACGATTCATCATCTCCAGAGACAATACCAAGAAGACCCTGTACCTGCAAATGAGC
AGTCTGAGGTCTGAGGACACAGCCTTGTATTACTGTGCACAGTGTTGTAACCACATCACG
AGTGTGTCAGAAACCCTGGAGAGCAGAAAGCTGCACTGCGACTGAGATGACAGAAGGATT
AATCCTTAGACTTGCTCAGAATTTGTAATTTTGAATGTCCATTCATTACCTCCTCCTCAG
AGTCCTATAGTGCCTTTGTCAGCTTTGTAAATGTCCATCGGTGAGTAAAGTGAAGATATT
TGGATAAACCTAAAATTCTTCACACTTTCTGAATCTATTCACCAGTGACCACCTCC

>M02293:67:000000000-LFYL9:1:1105:5825:8009 1:N:0:0 ATGTCA reverse complement 5-2 by 1- 1 insertion
GGCCGATTCACCATCTCCAGAGACAATACCAAGAAGACCCTGTACCTGCAAATGAGCAGT
CTGAGGTCTGAGGACACAGCCTTGTATTACTGTGCCACAGTGTTGTAACCACATCCTGAG
T

>M02293:67:000000000-LFYL9:1:1118:4543:16029 1:N:0:0 ATGTCA reverse complement 5-2 by 1- 1 insertion
GGCCGATTCACCATCTCCAGAGACAATACCAAGAAGACCCTGTACCTGCAAATGAGCAGT
CTGAGGTCTGAGGACACAGCCTTGTATTACTGTGTCACAGTGCTA

>M02293:67:000000000-LFYL9:1:2111:18651:8400 1:N:0:0 ATGTCA reverse complement 5-4 by 3-1/2(PG) 2 insertions
GGGCGATTCACCATCTCCAGAGACAATGCCAAGAACAACCTGTACCTGCAAATGAGCCAT
CTGAAGTCTGAGGACACAGCCATGTATTACTGTGGTCACAGTGTGGAGTCTTCACTGTGA
GCCCAGACAAAAACCTCCTTGCAGAGCAGCACTGCACCAACAGGGGGCATGGAGCATACA
CCAATAATGGGAAATCTACTTTAAA

>M02293:67:000000000-LFYL9:1:1110:18852:9656 1:N:0:0 ATGTCA reverse complement 5-12 by 2-6*03(by blast) 3 insertions
GGGCGATTCACCATCTCCAGAGACAATGCCAAGAACACCCTGTACCTGCAAATGAGCCGT
CTGAAGTCTGAGGACACAGCCATGTATTACTGTGAAGCACAGTGTGGGAAGTCCAATGTG
AGCCTGCACAAATACTTCTCTGCAGGGATGCTCACAACCAGCAGGGGGCGCTGAGGACCC
AAAGGGACTTCCCAGGATCTCTTCTGGAATCTAGGGAGCTCTGACCTGTGTCTATCAGCA
TGTGTTTCAATGTTAGAGTTCTTAGTTTTCCTTCCAGCAACAGAGATATTTTAGAGCCC

>M02293:67:000000000-LFYL9:1:1119:19064:19452 1:N:0:0 ATGTCA reverse complement 5-6 by 5-9 3 insertions
GGGCGATTCACCATCTCCAGAGACAATGCCAAGAACACCCTGTACCTGCAAATGAGCAGT
CTGAAGTCTGAGGACACAGCCATGTATTACTGTGGGTCACAGTGAGTGAATGTTACTGTG
AGCTCAAACTAAAACCTCCTGAAGAGCACCCAGGACCAGCAGGGGGCTGAGAGAGCACAG
TAACTTG

**SP**

>M02293:67:000000000-LFYL9:1:1101:21037:3627 1:N:0:0 CCGTCC reverse complement 5-2 by 1-19 0 insertions
GAGAGACGATTCATCATCTCCAGAGACAATACCAAGAAGACCCTGTACCTGCAAATGAGC
AGTCTGAGGTCTGAGGACACAGCCTTGTATTACTGTGCACAGTGCTACAAACACATCCTG
AGTGTGTCAGAAAACTTGGAGGTGCAGCAAGCTCCCTTGGGACTGACAAGGCTTAGAGAA
GGGCCGCTTGCAGATTTGCTT

>M02293:67:000000000-LFYL9:1:1118:16675:6391 1:N:0:0 CCGTCC reverse complement 5-4 by 1-19/36 0 insertions
GGGCGATTCACCATCTCCAGAGACAGTGCCAAGAACAACCTGTACCTGCAAATGAGCCAT
CTGAAGTCTGAGGACACAGCCATGTATTACTGTGCACAGTGCTACAAACACATCCTGAGT
GTGTCAGAAAACTTGGAGGTGCAGCAAGCTCCCTTGGGACTGAC

**Sort 3: RCseq, Aviti Sequencing, S. Andrews search; and shell-based searches for back to back RSS with up to 3 insertions. Pro-B pool (DEBC), Pro-B subsets (LN, LP, SN, SP) and large pre-B pool (Cp)**

**DEBC andrews**

>AV240405:AV_A_HW6069_PE300_200624:2409673462:1:10504:1440:0212 reverse complement 5-2 by 1-39 4 insertions
GAGAGACGATTCATCATCTCCAGAGACAATACCAAGAAGACCCTGTACCTGCAAATGAGC
AGTCTGAGGTCTGAGGACACAGCCTTGTATTACTGTGCAGGCACAGTGTTGTAACCACAT
CCTGAGTGTGTCAGAAAACCTGGAGGTGCAGCAAGCTCCCTTAGAACTGACAAGACTTAG
AAAAGTGTTGCTTGTAGATTTGCTTAGATGCAGTCATTTGAATAGTGTGTTTTTGTGTCT
ATTTAGTAAAATCCTATTGTGCTTTTTCAGCTTTGTAGAAGGATATCCATGAATTGCATG

CAGTTTACTAGGATGTCCCTAGAGTACCCCATGCCTTGTCAATATGGATAACAGTGAA

>AV240405:AV_A_HW6069_PE300_200624:2409673462:1:10104:0426:1067
reverse complement 5-2 by 12-3 0 insertions
GAGAGACGATTCATCATCTCCAGAGACAATACCAAGAAGACCCTGTACCTGCAAATGAGC
AGTCTGAGGTCTGAGGACACAGCCTTGTATTACTGTGCACAATGAGAAGATTCCAATGTC
AACCCACACACAAACCTCACTGCAGAGGGGATTTCAACCAGCAGGTGGTGCTGTTCGATG
TAAATGATCAGTAATTTCAAACATCTGTAAAAGAAAAAGAGAATGTCTTTGGCCAGGGCT
AAGTCTTTTCTGTATATGATTATAGGAACATCAGTTGTTCATGAATATCTTTTGGAGAAC
TAAATGACTATAGTGTTTCAATTACCGAG

>AV240405:AV_A_HW6069_PE300_200624:2409673462:1:11003:4897:0161
reverse complement 2-6-8 by 1-26 1 insertion
TCAAATCCAGACTGAGCATCAGCAAGGACAACTCCAAGAGCCAAGTTTTCTTAAAAATGA
ACAGTCTGCAAACTGATGACACAGCCATGTACTACTGTGCCACAGTGCTACAAACACATC
CTGAGTGTGTCAGAAACCCTGGAGGTGCAGCAAGCTCCCTTGGGATTGACAAGACTTAGA
GAATAGCCGCTTGCAGACTTCCTTAGATGCAGTCATTTGAATAGTGTGTTTTTGTGTCTA
TTTCTTAAAGTCCTATTGTGCTTTTTCTGCTTTTCAGAAGAAAATCAATGAATTGCATGC
AGTTTACTAGGATGTCCCTGGAGTACCCCATGCCTTGTCAATATGGATAACAGTGAACGC
TTTCATTGACATTTTTTCTTCTTTCATAAAACCACAC

>AV240405:AV_A_HW6069_PE300_200624:2409673462:1:10204:4020:1534
reverse complement 2-3 by 1-7 0 insertions
TCAAATCCAGACTGAGCATCAGCAAGGATAACTCCAAGAGCCAAGTTTTCTTAAAACTGA
ACAGTCTGCAAACTGATGACACAGCCACGTACTACTGTGCACAGTGGTGCAACCACATCC
CGACTGTGTCAGAAACCCTAGCAGAACAGGAAGCTTCCCTGGGACTGAGAATTCAGAAAA
GACTAACCTGTAGGCTTGATGAAAAATAATCATTTTGGGCACTAATTTTTATGCATGCTC
CTTACACTCTTACAGCGCCTTTTTCAACTATGTAAATA

>AV240405:AV_A_HW6069_PE300_200624:2409673462:1:11301:0028:1956
reverse complement 5-2 by 1-76 0 insertions
GAGAGACGATTCATCATCTCCAGAGACAATACCAAGAAGACCCTGTACCTGCAAATGAGC
AGTCTGAGGTCTGAGGACACAGCCTTGTATTACTGTGCACAGTGTTACAACCACATCCTG
AGTGTGTCAGAAACCCTGTAGGAGCAGGAAGCTGCACTGAGACTGAGATGACAGAAAGAT
TAATCTTTAGACTTGCTCAGAAATTGTAATTTTGAATGTCCATTTATTACCTCCTACTAA
CAGTCGTATAGTATCTTTGTCAACTTTGTCATCGTTTTCTATGAATAAAGTAAGGCTGTT
TGGATAAACTATCAATTCACAATACCTTGTGAATCTCTTCACCATTGA

>AV240405:AV_A_HW6069_PE300_200624:2409673462:1:11202:1313:3316
reverse complement 5-6 by 1-81 0 insertions
GGCCGATTCACCATCTCCAGAGACAATGCCAAGAACACCCTGTACCTGCAAATGAGCAGT
CTGAAGTCTGAGGACACAGCCATGTATTACTGTGCACAGTGTTGTAACCACATCCTGAGT
GTGTCAGAAAACCTGGGGCAGAAGAAAGATAAGCTGGGACTGAGAAGACAGAAAAATTAA
TCCTTAGATTTGCTCAGAAATCATAGTTTTGAATGCCTATTTATTTCCTTCTCCTCACAG
ATCTATAGTGCTTTTGTCAGCTTCTTAAATGTCCATCTATGAGT

>AV240405:AV_A_HW6069_PE300_200624:2409673462:1:10705:0404:2814
reverse complement 2-4 by 1-82 3 insertions
TCATATCCAGACTGAGCATCAGCAAGGACAACTCCAAGAGCCAAGTTTTCTTTAAAATGA
ACAGTCTGCAAGCTGATGACACTGCCATATACTACTATGTCCCACAGTGTTACAACCACA
TCCTGAGAGTGTCAGAAACCGTGGAGGAGCAGGAAGCTTCCCTGGGCCTGAGATGACAGA
AAGATTAATCTTTAGACTTGCTCAGAAATCATAATTACTTTGTAAACATCCATCTATGAA
TAAAGTGATGCTGTTTGGAGAAACCTACACCTTCTGAATCTCTTCACCTGTGACCTGTTT
CTTATTCAATAAAACAATAAAAAGCAAAAGTGTGTTTTTGCATGATAAAAATATTACTGA
ATCCTGAGTGACTAGAACTTTTTTAAAGTTGTTGGGAATAGTCTGTAATAGAAATTTGAT

CTATGTGCAAGCCACCAACATTTGGTCAAACAAACAATGCTTTATAGTCTTATGTAACTT
AGTGACCTCAAGTGTGTTAACTCTTGTCCCATGTTCTAGCAGAAATTCCTGTCTCTTAGA
AGGTG

>AV240405:AV_A_HW6069_PE300_200624:2409673462:1:12005:3725:1678
reverse complement 5-2 by 1-36 0 insertion
GGGCGATTCACCATCTCCAGAGACAATACCAAGAAGACCCTGTACCTGCAAATGAGCAGT
CTGAGGTCTGAGGACACAGCCTTGTATTACTGTGCACAGTGCTACAAACACATCCTGAGT
GTGTCAGAAAACTTGGAGGTGCAGCAAGCTCCCTTGGGACTGACAAGACTTAGAGAAGTG
CCGCTTGCAGATTTGATTAAATGTAATCATTTGAATAGTGTGTTTTTGTGTCTATTTCTT
AAAGACCTATTTTTCTTTTTTAGCTTTGCACAAGGACATCCATGAATTACATGCAGTTTA
CTAGGATGTCCCTGGAGTACCCCCATGCCTTGTCAATATGGATAACAGTGAACACTTTCC
TTGAAAAATTTCTTCTTTCA

**DEBC shell**
>AV240405:AV_A_HW6069_PE300_200624:2409673462:1:21605:4021:3544
5-2 by 3-6  0 insertions
GAGAGACGATTCATCATCTCCAGAGACAATACCAAGAAGACCCTGTACCTGCAAATGAGC
AGTCTGAGGTCTGAGGACACAGCCTTGTATTACTGTGCACAGTGTGGAGTCTTCACTGTG
AGCCCAGACATAAACCTCCTGTAGAGCAGCTCTGTACCAACAGGGGGCTTGTCACATACA
CTTAGCACAGGAAATCTATTTTAGGGTATGTTAAACACAATTCTTGCCCAAAACTCTGGC
AGGAGACTCCTGCTGAAAAGGGAGACACATTAGTGA

>AV240405:AV_A_HW6069_PE300_200624:2409673462:1:21001:1581:2018
5-4 by 13-1 Pseudogene  0 insertions
GGGCGATTCACCATCTCCAGAGACAATGCCAAGAACAACCTGTACCTGCAAATGAGCCAT
CTGAAGTCTGAGGACACAGCCATGTATTACTGTGCACAGTGTGGAGTCTTCAGTGTAGGC
CCAGACATAAACCTCCTTGTAGAGCAGCTCT

>AV240405:AV_A_HW6069_PE300_200624:2409673462:1:21403:1990:1232
1-11 by 1-84 (not in VH89, low recomb)  0 insertions
AGTTCAAGGGCAAGGCCACATTCTCTGTAGACCGGTCCTCCAGCACAGTGTACATGGTGT
TGAACAGCCTGACATCTGAGGACCCTGCTGTCTATTACTGTGCACAGTGTTACAACCACA
TCCTGAGTGTGTCAGAAACCCTGGATGAGCAGGAAGCTGCACTGCGACTGAGATGACAGA
AAGATTAATCCTTAGACTTGCTCAGAAATTGTAATTTTGAATGTCCATTTATTACCTCCT
CCTCAGAGTCCTATAGTGCCTTTGTCAGCTTTGCAAACGTTCATCTATGAATAATGTCAT
TCTGTTTGGATAAACCTACAGTTCACCATACCTTGTGAATTTCTTCATCCGTGACCACTT
TCCATAAAAAAGAACAAATTTTATTCAATAGAACAAAAAAAGTGAAAAGTATGTTTATAC
ATGATAAAAATATTACTGGATCCTGAGTGATCAG

**LP Andrews**
>AV240405:AV_A_HW6069_PE300_200624:2409673462:1:20103:2233:0449
reverse complement 2-3 by 1-82 0 insertions
TCAAATCCAGACTGAGCATCAGCAAGGATAACTCCAAGAGCCAAGTTTTCTTAAAACTGA
ACAGTCTGCAAACTGATGACACAGCCACGTACTACTGTGCACAGTGTTACAACCACATCC
TGAGAGTGTCAGAAACCGTGGAGGAGCAGGAAGCTTCCCTGGGCCTGAGATGACAGAAAG
ATTAATCTTTAGACTTGCTCAGAAATCATAATTACTTTGTAAACAT

>AV240405:AV_A_HW6069_PE300_200624:2409673462:1:12102:1974:0559
reverse complement 1-11 by 1-19 0 insertions
CATGGGCAAGGCCACATTCTCTGTAGACCGGTCCTCCAGCACAGTGTACATGGTGTTGAA
CAGCCTGACATCTGAGGACCCTGCTGTCTATTACTGTGCACAGTGCTACAAACACATCCT
GAGTGTGTCAGAAAACTTGGAGGTGCAGCAAGCTCCCTTGGGACTGACAAGGCTTAGAGA
AGGGCCGCTTGCAGATTTGCTTAAATGTAATCATTTGAATACTGTGTTTTTGTGTCTATT
TCTTAAAGACCTATTTTTCTTTTTTAGCTTTGCACAAGGCCATCCATGAATTACATGCAG

TTTACTAGGATGTCCCTGGAGTACCCCCATGCCTTGTCAATATGGATAACAGTGAACACT
TTCCTTGAAAAAATTTCTTCTTTCATAAAACAACAAAAAAATGGAAAATGTGATT

**LP shell**
>AV240405:AV_A_HW6069_PE300_200624:2409673462:1:10803:0664:0432
5-4 by 1- 3 insertions
GGGCGATTCACCATCTCCAGAGACAATGCCAAGAACAACCTGTACCTGCAAATGAGCCAT
CTGAAGTCTGAGGACACAGCCATGTATTACTGTGGGCCACAGTGTTGTAACC

**SN Andrews**
>AV240405:AV_A_HW6069_PE300_200624:2409673462:1:11704:2545:2656
reverse complement 5-6 by 9-4 2-insertions
GGGCGATTCACCATCTCCAGAGACAATGCCAAGAACACCCTGTACCTGCAAATGAGCAGT
CTGAAGTCTGAGGACACAGCCATGTATTACTGTGCCCACAGTGTGAAAACCACATCCTGA
GTGTGTCAGAAACCATGAGGAGAAGGTGGTTCAGCTAAGCCCAGACAAAAAGTGAGAAAA
CATTCTCTCCTTCATTATGGACCACAAATACGAGCTTACTGACAATAGATAGAAAATTCA
CATATGGTGAGCCTCAGAATGTTCTCAGTGGGTTGTGACAGACTACCTTAATGAAAGGAG
AAGGGAGTTTAGGGATCAGGTGGGAAGGGGGATGGGGGCATTCA

>AV240405:AV_A_HW6069_PE300_200624:2409673462:1:21402:4565:2313
reverse complement 1-39 by 1-75 2 insertions altered RSS
AGTTCAAGGGCAAGGCCACATTGACTGTAGACCAATCTTCCAGCACAGCCTACATGCAGC
TCAACAGCCTGACATCTGAGGACTCTGCAGTCTATTACTGTGCCTTCAGTGTTACAACCA
CATCCTGAGTGTGTCAGAAACCCTGGAGGAGCAGGAAGCTGCACTGGGACTGAGATGACA
GAAAGATTAATCCTTAGACTTTCTCAGAATTTTAATTCTGAATGTCCATTTATTACCTCC
TCCTCAGAGTCCTATAGTGCCTTTGTCAGCTTTGTAAATGTCCATCTATGAGTAAAGTGA
T

>AV240405:AV_A_HW6069_PE300_200624:2409673462:1:10104:3147:0858
reverse complement 1-11 by 8-12 4 insertions altered RSS
CATGGGCAAGGCCACATTCTCTGTAGACCGGTCCTCCAGCACAGTGTACATGGTGTTGAA
CAGCCTGACATCTGAGGACCCTGCTGTCTATTACTGTGGAATAATTTTGATACAGCTTAA
GTTTTCAGCTGTACAGTATTTCAGGCTGGTGTTCAGCTCTGTTCCTTCCAATTATATCAT
AAGAGGGTATGGCCGCAGTTTTCCTCTTGCCTCTGTGGTTTCCTGTTTGCTCTTAGGTTA
AGAGCTTTCTCTTCCTGTTATTCATCC

>AV240405:AV_A_HW6069_PE300_200624:2409673462:1:10104:2610:3463
reverse complement 5-2 by 1-77 0 insertions
GAGAGACGATTCATCATCTCCAGAGACAATACCAAGAAGACCCTGTACCTGCAAATGAGC
AGTCTGAGGTCTGAGGACACAGCCTTGTATTACTGTGCACAGTGTTGTAACCACATCCTG
AGTGTGTCAGAAACACTGGAGGAGCAGGAAGCTGCACTGGGACTGAGATGACAGAAAGAT
TAATCCTTAGACTTGCTCAGAAATTGTAATTTTGAATGTCCATGTATTG

>AV240405:AV_A_HW6069_PE300_200624:2409673462:1:12004:0643:2327
reverse complements 5-4 by 1-26 0 insertions
GGGCGATTCACCATCTCCAGAGACAATGCCAAGAACAACCTGTACCTGCAAATGAGCCAT
CTGAAGTCTGAGGACACAGCCATGTATTACTGTGCACAGTGCTACAAACACATCCTGAGT
GTGTCAGAAACCCTGGAGGTGCAGCAAGCTCCCTTGGGATTGACAAGACTTAGAGAATAG
CCGCTTGCAGACTTCCTTAGATGCAGTCATTTGAATAGTGTGTTTTTGTGTCTATTTCTT
AAAGTCCTATTGTGCTTTTTCTGCTTTTCAGAAGAAAATCAATGAATTGCATGCAGTTTA
CTAGGATGTCCCTGGAGTACCCCATGCCTTGTCAATATGGATAACAGTGAACGCTTTCAT
TGACATTTTTTCTTCT

>AV240405:AV_A_HW6069_PE300_200624:2409673462:1:12004:4401:3484
reverse complement 5-2 by 1-82 0 insertions
GAGAGACGATTCATCATCTCCAGAGACAATACCAAGAAGACCCTGTACCTGCAAATGAGC
AGTCTGAGGTCTGAGGACACAGCCTTGTATTACTGTGCACAGTGTTACAACCACATCCTG
AGAGTGTCAGAAACCGTGGAGGAGCAGGAAGCTTCCCTGGGCCTGAGATGACAGAAAGAT
TAATCTTTAGACTTGCTCAGAAATCATAATTACTTTGTAAACATCCATCTATGAATAAAG
TGATGCTGTTTGGAGAAACCTACACCTTCTGAATCTCTTCACCTGTGACCTGTTTCTTAT
TCAATAAAACAATAAAAAGCAAAAGTGTGTTTTTGCATGATAAAAAAATTACTGAATCCT
GAGTGACTAGAACTTTTTTAAAGTTGTTGGGAATAGTCTGTAATAGAAATTTGATCTATG
TGCAAGCCACCAACATTTGGTCAAACAAACAATGCTTTATAGTCTTATGTA

>AV240405:AV_A_HW6069_PE300_200624:2409673462:1:10203:5301:2116
reverse complement 3-1 by 1-69 4 insertions all but 1 of 3-1 3’
CCCTCAAAAGTCGAATCTCCATCACTCATGACACATCTAAGAACCATTTCTTCCTGAAGT
TGAATTCTGTGACTACTGAGGACACAGCCACATATTACTGTGCAAGAGGAGTGTTGCAAC
CACATCCTGAGAGTGTCAGAAACCCTGGAGGAGTAGCAAACTGCCCTGAGACTGAGGAGA
CTCAGAGAAGGTTTGCTTGTAGACTTGCTCAGATACAGCCAGGATGGTGTGTAGTATGGG
CCCCAGACATGTA

>AV240405:AV_A_HW6069_PE300_200624:2409673462:1:20401:3658:2741
reverse complement 2-2 by 1-7 0 insertions
TCATATCCAGACTGAGCATCAGCAAGGACAATTCCAAGAGCCAAGTTTTCTTTAAAATGA
ACAGTCTGCAAGCTGATGACACAGCCATATATTACTGTGCACAGTGGTGCAACCACATCC
CGACTGTGTCAGAAACCCTAGCAGAACAGGAAGCTTCCCTGGGACTGAGAATTCAGAAAA
GACTAACCTGTAG

>AV240405:AV_A_HW6069_PE300_200624:2409673462:1:22101:4882:1934
reverse complement 2-4 by 5-12 0 insertions
TCATATCCAGACTGAGCATCAGCAAGGACAACTCCAAGAGCCAAGTTTTCTTTAAAATGA
ACAGTCTGCAAGCTGATGACACTGCCATATACTACTGTGCACAATGAGGAAATGTTACTG
TGAGCTCAAACTAAAACCTCCTGCAGAGCACCCAGGACCAGCAGGGGGCGCAGAGAGCAC
ATGGAGTTCTGATTCACAGAAGAGTTACAGCCTGTACAATTAGACCCAATCTTCAACAAA
CCGTCAAAATATTCGATCCAAAATTGTTCCTGTCTAAAAGTAATTCAAGGACAAAATGGA
CCAGAGACTGAAGAAATGGCTGACCTGTGACCCTCCCAACTTTGGATCTATCTCATAGGC
AGGTACCAAACCTTGACATTTGTCACTGACACTGTATTGTGCTTGCAGACAGGAGCATAG
CATGGCTGACCTCTAAGAGGCTCTGCAAGCACCTGAATGAGACAGATG

**SP andrews**
>AV240405:AV_A_HW6069_PE300_200624:2409673462:1:20102:0907:2368
reverse complement 5-2 by 1-55 0 insertions
GAGAGACGATTCATCATCTCCAGAGACAATACCAAGAAGACCCTGTACCTGCAAATGAGC
AGTCTGAGGTCTGAGGACACAGCCTTGTATTACTGTGCACAGTGTTGCAACCACATCCTG
AGAGTGTCCAAAACCCTGGAGGAGTAGCAAACTGCCCTGGGACTGTGGAGACTCAGAGAA
GTTTTGCTTGTAGACTTGCTCAGATACAGCCATTTAGATAGTCCAATTTTGTGTTTGTTA
CTTAGAGTTCTGTAGTGCTTTTGTCCACTTTGCAAATGGACATCGATTAATCATATGATA
TTGATTTAGGATGCTTTGAGAGTATGCCATACCTTGTCAATTACCTTCTTCACCAGTGAC
CATCTCTGTTGACATATTTCTTTTGTA

>AV240405:AV_A_HW6069_PE300_200624:2409673462:1:10103:3941:0457
reverse complement 3-1 by 15-2 0 insertions
CCCTCAAAAGTCGAATCTCCATCACTCATGACACATCTAAGAACCATTTCTTCCTGAAGT
TGAATTCTGTGACTACTGAGGACACAGCCACATATTACTGTGCACAGTGTAACAGCTCAT

ATCTGAAGCATGTCAAAAAGTCTGAAGGCAGGAAGCTATTTTGGGTCTGATATTACCACA
AAGATTAACCAAAGCAGTTGCTCAAAGCCTGTGGTTAAATGTTGAA

>AV240405:AV_A_HW6069_PE300_200624:2409673462:1:10104:4315:0576
reverse complement 5-2 by 1-9 0 insertions
GAGAGACGATTCATCATCTCCAGAGACAATACCAAGAAGACCCTGTACCTGCAAATGAGC
AGTCTGAGGTCTGAGGACACAGCCTTGTATTACTGTGCACAGTGTTGTAACCACATCCTG
AGTGTGTCAGAAACTCTGGAGGAGCAGCAAGCTGCCCTGGGTCTGAAATATCAGAAAAGG
CTAACATTTAGACTTCCTCAGAAATAACCATTTTGAGTGCCTATTTTTCGTTTGTTAGTT
TCCTACACACAGACTTATAGTGCCTTTGTCAAC

>AV240405:AV_A_HW6069_PE300_200624:2409673462:1:20104:0057:3384
reverse complement 5-2 by 12-3 0 insertions
GAGAGACGATTCATCATCTCCAGAGACAATACCAAGAAGACCCTGTACCTGCAAATGAGC
AGTCTGAGGTCTGAGGACACAGCCTTGTATTACTGTGCACAATGAGAAGATTCCAATGTC
AACCCACACACAAACCTCACTGCAGAGGGGATTTCAACCAGCAGGTGGTGCTGTTCGATG
TAAATGATCAGTAATTTCAAACATCTGTAAAAGAAAAAGAGAATGTCTTTGGCCAGGGCT
AAGTCTTTTCTGTATATGATTATAGGAACATCAGTTG

>AV240405:AV_A_HW6069_PE300_200624:2409673462:1:20601:2543:1099
reverse complement 2-3 by 1-9 0 insertions
TCAAATCCAGACTGAGCATCAGCAAGGATAACTCCAAGAGCCAAGTTTTCTTAAAACTGA
ACAGTCTGCAAACTGATGACACAGCCACGTACTACTGTGCACAGTGTTGTAACCACATCC
TGAGTGTGTCAGAAACTCTGGAGGAGCAGCAAGCTGCCCTGGGTCTGAAATATCAGAAAA
GGCTAACATTTAGACTTCCTCAGAAATAACCATTTTGAGTGCCTATTTTTCGTTTGTTAG
TTTCCTACACACAGACTTATAGTGCCTTTGTCAACTTTGCATAAG

>AV240405:AV_A_HW6069_PE300_200624:2409673462:1:10301:3991:1183
reverse complement 5-2 by 6-3 3 insertions
GAGAGACGATTCATCATCTCCAGAGACAATACCAAGAAGACCCTGTACCTGCAAATGAGC
AGTCTGAGGTCTGAGGACACAGCCTTGTATTACTGTGAGACACAGTGAGAAGTCTTCATT
GTGAGTCTAGACACAAACTTACCCAAAGGAGCTCTCAGTACCAGCAGGGGGAGCACAGTG
ACAATCGAATCCATAAATGGGCTATTGTTTACAGGGATCTGGGCAGGTGAGACCACTTTC
TCAGTAGTCCCTTTCCTTCCTCCACCATCTGGAG

**SP shell**
>AV240405:AV_A_HW6069_PE300_200624:2409673462:1:21405:0623:2734
5-2 by 8-7 Pseudogene  0 insertions
GAGAGACGATTCATCATCTCCAGAGACAATACCAAGAAGACCCTGTACCTGCAAATGAGC
AGTCTGAGGTCTGAGGACACAGCCTTGTATTACTGTGCACATTGACACAGCCTCAGTTTT
CATCTGTACATTATTTCAGGCTGATTCTCAGTTGTGTTTCTTTCTTTTATATTACAAGAG
GGTATGGCCACAGATTTCCTCTGACCTCTGTGGTTTCCTGTTTGCTCTTAGGTCAAGAGC
CCTCTTCCTGTTATTCACTGTAGGTCCCTATTAATGTGTTCATAATTGATTCTGTCTGTG
TTTTGCTTTTAAAGCCGATGGAGGTTTGGTGCCTACACCAGGTGGGTCAGAGTTATAGTT
TCACTCAGTTTTAGACTTCAAATTCCCACAGTAGAATACTCCAAATTATTGTAATTTTCA
ACTTCAGTAAACAGCTGAAGATGGGAATTACTGGATGTGTATTCTCAAATTGAGTCATTG
AAAAAAAGAGTGTTC

**Cp Andrews**
>AV240405:AV_A_HW6069_PE300_200624:2409673462:1:11503:0524:0529
reverse complement 5-4 by 9-2 2 insertions
GGCCGATTCACCATCTCCAGAGACAATGCCAAGAACAACCTGTACCTGCAAATGAGCCAT
CTGAAGTCTGAGGACACAGCCATGTATTACTGTGCGCACAGTGTGAAAACCACATCCTGA
GGGTGTCAGAAACCATGAGGAGAAGGTGGTTCAGCTGTGTCCAGAAGCAACCAGAGGAAA
CATTCTCTCCTTGATGTTTGGCCACAATTATGAGATTACTGACAACACATATAATAGTCA

TATATGGTCAGCCACAGAATGTTCTCAGTGGGATTGTGACAGATCAGATTGATGAAAGGA
AGAGGGACCGTTTCATAGCATCTTTAATTTGGTATATATAAACTGTTAATATGAAAATAT
CA

>AV240405:AV_A_HW6069_PE300_200624:2409673462:1:21403:1064:3470
reverse complement 2-2 by 1-26 0 insertions
TCATATCCAGACTGAGCATCAGCAAGGACAATTCCAAGAGCCAAGTTTTCTTTAAAATGA
ACAGTCTGCAAGCTGATGACACAGCCATATATTACTGTGCACAGTGCTACAAACACATCC
TGAGTGTGTCAGAAACCCTGGAGGTGCAGCAAGCTCCCTTGGGATTGACAAGACTTAGAG
AATAGCCGCTTGCAGACTTCCTTAGATGCAGTCATTTGAATAGTGTGTTTTTGTGTCTAT
TTCTTAAAGTCCTATTGTGCTTTTTCTGCTTTTCAGAAGAAAATCAATGAATTGCATGCA
GTTTACTAGGATGTCCCTGGAGTACCCCATGCCTTGTCAATATGGATAACAGTGAACGCT
TTCATTGACATTTTTTCTTCTTTCATAAAACCACACAAAAGGGAAAATGTGATTATGCAT
GATAAAAGTATGTTTTTATTGTGTAAAAGACTTGAACTTTGGAATAGGTGGGAATGAATT
ATCATAGGAATTTGACCCGTAAAATTC

>AV240405:AV_A_HW6069_PE300_200624:2409673462:1:10201:3934:2129
reverse complement 2-2 by 1-19 0 insertions
TCAAATCCAGACTGAGCATCAGCAAGGACAATTCCAAGAGCCAAGTTTTCTTTAAAATGA
ACAGTCTGCAAGCTGATGACACAGCCATATATTACTGTGCACAGTGCTACAAACACATCC
TGAGTGTGTCAGAAAACTTGGAGGTGCAGCAAGCTCCCTTGGGACTGACAAGGCTTAGAG
AAGGGCCGCTTGCAGATTTGCTTAAATGTA

>AV240405:AV_A_HW6069_PE300_200624:2409673462:1:10703:5212:2446
reverse complement 2-2 by 1-22 0 insertions
TCAAATCCAGACTGAGCATCAGCAAGGACAATTCCAAGAGCCAAGTTTTCTTTAAAATGA
ACAGTCTGCAAGCTGATGACACAGCCATATATTACTGTGCACAGTGCTACAAACACATCC
TCAGTGTGTCAGAAATCCTGGAGGTGCAGCAAGCTCCCTTGGGACTGACAAGACTTAGAG
AATAGTTGCTTGCAGACGTGCTTAGGTGCAGACATTTGGATAGTGTGTTTTTGTGTCTAT
TTCTTAAAGACCTAGTATGCTTTTTCAGCTTTTCAGAAGAAAATCAAT

>AV240405:AV_A_HW6069_PE300_200624:2409673462:1:20802:4607:3201
reverse complement 5-2 by 1-39 0 insertions
GAGAGACGATTCATCATCTCCAGAGACAATACCAAGAAGACCCTGTACCTGCAAATGAGC
AGTCTGAGGTCTGAGGACACAGCCTTGTATTACTGTGCACAGTGTTGTAACCACATCCTG
AGTGTGTCAGAAAACCTGGAGGTGCAGCAAGCTCCCTTAGAACTGACAAGACTTAGAAAA
GTGTTGCTTGTAGATTTGCTTAGATGCAGTCATTTGAATAGTGTGTTTTTGTGTCTATTT
AGTAAAATCCTATTGTGCTTTTTCAGCTTTGTAGAAGGATATCCATGAATTGCATGCAGT
TTACTAGGATGTCCCTAGAGTACCCCATGCCTTGTCAATATGGATAACAGTGAACACTTT
CATTGACTTATTTCTTCTT

>AV240405:AV_A_HW6069_PE300_200624:2409673462:1:10404:2516:0548
reverse complement 5-2 by 14-1 0 insertions
GAGAGACGATTCATCATCTCCAGAGACAATACCAAGAAGACCCTGTACCTGCAAATGAGC
AGTCTGAGGTCTGAGGACACAGCCTTGTATTACTGTGCACAGTCTTGCAAACACATCCTG
AGAGTGTCATAAACCATAAAGTGCAGGAAGCTGCCTGGAACTGAGATGACAGACTATACT
ACCCTGAAGAAATAGCCATTTTGAGTGTCCCAATTTCTGCCTCCTCCTCA

>AV240405:AV_A_HW6069_PE300_200624:2409673462:1:20401:1492:3626
reverse complement 2-2 by 9-3 4 insertions
TCATATCCAGACTGAGCATCAGCAAGGACAATTCCAAGAGCCAAGTTTTCTTTAAAATGA
ACAGTCTGCAAGCTGATGACACAGCCATATATTACTGTGATCTCACAGTGTGAAAACCAC
ATCCTGAGGGTGTCAAAAACCATGAGGAGAAGGTGGTTCAGCTGTGTCCAGAAGCAACCA
GAGGAAACATTCTCTCCTTGGTGTTTGGCCACAATTATGAGATTACTGACAACACATATA
ATAGTCA

>AV240405:AV_A_HW6069_PE300_200624:2409673462:1:10604:0989:1981
reverse complement 5-2 by 1-19 3 insertions
GAGAGACGATTCATCATCTCCAGAGACAATACCAAGAAGACCCTGTACCTGCAAATGAGC
AGTCTGAGGTCTGAGGACACAGCCTTGTATTACTGTGGCCCACAGTGCTACAAACACATC
CTGAGTGTGTCAGAAAACTTGGAGGTGCAGCAAGCTCCCTTGGGACTGACAAGGCTTAGA
GAAGGGCCGCTTGCAGATTTGCTTAAATGTAATCATTTGAATACTGTGTTTTTGTGTCTA
TTTCTTAAAGACCTATTTTTCTTTT

**Cp shell**
>AV240405:AV_A_HW6069_PE300_200624:2409673462:1:12104:0250:1332
5-2 by 8-2 (ORF) 0 insertions
GAGAGACGATTCATCATCTCCAGAGACAATACCAAGAAGACCCTGTACCTGCAAATGAGC
AGTCTGAGGTCTGAGGACACAGCCTTGTATTACTGTGCACAGTGGTGCAACCGTGACCCA
CAGCTGTGCAATATTTCAGGATGGATCTCATTTCTGTTCCTTCCTAATAGAGAGGAGGAG
TTGGAGCGGGTCCACATTTCCTGCTAGCCTCTGGGGTTTCCTTTTTACTGTGAGCTTCAG
AGGCTAGGCTTCCTGTTACGCCCCCTAAGTCATTTTTCAAACCTTTTTGTTGTTTCTTTA
TAGATTTGACATCATGAACCCCATTTTTACTCATCTTCCCCTCTCCCACACACCCACTCT
CCAACCTTGGAACCTCCTCCCCAA

>AV240405:AV_A_HW6069_PE300_200624:2409673462:1:20203:3292:1480
5-12 by 2-6*03 0 insertions
GGCCGATTCACCATCTCCAGAGACAATGCCAAGAACACCCTGTACATGCAAATGAGCCGT
CTGAAGTCTGAGGACACAGCCATGTATTACTGTGCACAGTGTGGGAAGTCCAATGTGAGC
CTGCACAAATACT

>AV240405:AV_A_HW6069_PE300_200624:2409673462:1:10603:4885:0698
5-2 by 1-18 to 36, 0 insertions
GAGAGACGATTCATCATCTCCAGAGACAATACCAAGAAGACCCTGTACCTGCAAATGAGC
AGTCTGAGGTCTGAGGACACAGCCTTGTATTACTGTGCACAGTGCTACAAACA
